# Brain/MINDS Beyond Human Brain MRI Project: A Protocol for Multi-Site Harmonization across Brain Disorders Throughout the Lifespan

**DOI:** 10.1101/2020.05.05.076273

**Authors:** Shinsuke Koike, Saori C Tanaka, Tomohisa Okada, Toshihiko Aso, Michiko Asano, Norihide Maikusa, Kentaro Morita, Naohiro Okada, Masaki Fukunaga, Akiko Uematsu, Hiroki Togo, Atsushi Miyazaki, Katsutoshi Murata, Yuta Urushibata, Joonas Autio, Takayuki Ose, Junichiro Yoshimoto, Toshiyuki Araki, Matthew F Glasser, David C Van Essen, Megumi Maruyama, Norihiro Sadato, Mitsuo Kawato, Kiyoto Kasai, Yasumasa Okamoto, Takashi Hanakawa, Takuya Hayashi, Brain/MINDS Beyond Human Brain MRI Group

## Abstract

Psychiatric and neurological disorders are afflictions of the brain that can affect individuals throughout their lifespan. Many brain magnetic resonance imaging (MRI) studies have been conducted; however, imaging-based biomarkers are not yet well established for diagnostic and therapeutic use. This article describes an outline of the planned study, the Brain/MINDS Beyond human brain MRI project (FY2018 ∼ FY2023), which aims to establish clinically-relevant imaging biomarkers with multi-site harmonization by collecting data from healthy traveling subjects (TS) at 13 research sites. Collection of data in psychiatric and neurological disorders across the lifespan is also scheduled at 13 sites, whereas designing measurement procedures, developing and analyzing neuroimaging protocols, and databasing are done at three research sites. The Harmonization protocol (HARP) was established for five high-quality 3T scanners to obtain multimodal brain images including T1 and T2-weighted, resting state and task functional and diffusion-weighted MRI. Data are preprocessed and analyzed using approaches developed by the Human Connectome Project. Preliminary results in 30 TS demonstrated cortical thickness, myelin, functional connectivity measures are comparable across 5 scanners, providing high reproducibility and sensitivity to subject-specific connectome. A total of 75 TS, as well as patients with various psychiatric and neurological disorders, are scheduled to participate in the project, allowing a mixed model statistical harmonization. The HARP protocols are publicly available online, and all the imaging, demographic and clinical information, harmonizing database will also be made available by 2024. To the best of our knowledge, this is the first project to implement a rigorous, prospective harmonization protocol with multi-site TS data. It explores intractable brain disorders across the lifespan and may help to identify the disease-specific pathophysiology and imaging biomarkers for clinical practice.

## 1. Introduction

Psychiatric and neurological disorders are afflictions of the brain that can affect individuals throughout their lifespans. Using the disability-adjusted life years (DALYs), which is a measure of disease burden proposed by the World Health Organization Global Burden of Disease study, in 2010 mental and behavioral disorders accounted for 7.4% of the total DALYs and neurological disorders accounted for 3.0% (Murray et al., 2012), up from 5.4% and 1.9% in 1990, respectively. Since the 1990s, technical advances in magnetic resonance imaging (MRI) have allowed detailed analysis of the organization of brain function and structure in humans. Recent high-quality MRI studies with a large cohort are expected to provide neurobiological and life-span information in healthy subjects (Glasser et al., 2016b; Harms et al., 2018; Miller et al., 2016), which will hopefully provide diagnostic utility for patients with psychiatric and neurological disorders (Drysdale et al., 2017; Elliott et al., 2018b; Koutsouleris et al., 2015; Nunes et al., 2018). However, the diagnostic value of brain MRI in psychiatric disorders has not yet been established, presumably because effect sizes tend to be small and overlap with variability in healthy individuals (Yamashita et al., 2019). Protocols of scanning and analysis have rarely been standardized across projects, though that has begun to change - especially for large projects such as the Human Connectome Project (HCP; (Van Essen et al., 2012)), UK Biobank (Miller et al., 2016), and the Adolescent Brain Cognitive Development (ABCD) project (Casey et al., 2018).

### 1.1. Previous multi-site neuroimaging studies for neuropsychiatric disorders

Several brain imaging projects have attempted to identify suitable biomarkers in neuropsychiatric diseases. Recent multi-site neuroimaging mega studies have revealed well-replicated and clinically applicable findings from structural images; the Enhancing NeuroImaging Genetics through Meta-Analysis Consortium in the U.S. (n = 4,568) and the Cognitive Genetics Collaborative Research Organization in Japan (n = 2,564) replicated findings that patients with schizophrenia have volumetric alterations of subcortical structures when compared to healthy controls (Okada et al., 2016; van Erp et al., 2016). The findings were partly evident in other psychiatric disorders, such as bipolar disorder (BPD) and major depressive disorder (MDD) (Hibar et al., 2018; Schmaal et al., 2017; Schmaal et al., 2016; van Erp et al., 2016). Using resting-state functional MRI (rsfMRI), a multi-site study successfully developed generalized classifiers for psychiatric disorders. The Decoded Neurofeedback (DecNef) Project (https://bicr.atr.jp/decnefpro/), a multi-site neuroimaging study in Japan (12 sites, n = 2,409), developed a generalized classifier for autism spectrum disorder (ASD) with a high accuracy— not only for the data in three Japanese sites (85%) but also for the Autism Brain Imaging Data Exchange dataset (75%) (Yahata et al., 2016). The project also quantified the spectrum of psychiatric disorders by applying the ASD classifier to other multi-disorder datasets (schizophrenia, MDD, and attention-deficit/hyperactivity disorder). Therefore, the focus of mega-analyses is shifting from features found in case-control studies to cross-disease comparisons that can identify common and disease-specific features.

In the field of neurodegenerative disease, the Alzheimer’s Disease Neuroimaging Initiative (ADNI) is one of many major multi-site neuroimaging and biomarker studies of Alzheimer’s disease (AD) and mild cognitive impairment (MCI) that was started in 2005 in North America (Mueller et al., 2005; Weiner et al., 2015). It contributed to the development of blood and imaging biomarkers, the understanding of the biology and pathology of aging, and to date has resulted in over 1,800 publications. ADNI also impacted worldwide ADNI-like programs in many countries including Japan, Australia, Argentina, Taiwan, China, Korea, Europe, and Italy. The Japanese ADNI (J-ADNI) conducted a multi-site neuroimaging study on cognitively normal elderly patients, MCI, and mild AD (n = 537), which emphasized the harmonization of the protocol and procedures with the ADNI (Iwatsubo et al., 2018). J-ADNI also developed machine learning techniques using feature-ranking, a genetic algorithm, and a structural MRI-based atrophy measure to predict the conversion from MCI to AD (Beheshti et al., 2017). Inspired by the Parkinson’s Progression Markers Initiative (PPMI; (Parkinson Progression Marker Initiative, 2011), the Japanese (J-) PPMI team has also started a cohort in patients with rapid eye movement sleep behavioral disorder, which is regarded to be prodromal to Parkinson’s disease (PD) (Mukai and Murata, 2017).

These previous mega-studies have contributed to the discovery of potential mechanisms and biomarkers of multiple brain disorders. However, most of these imaging biomarkers have a relatively small effect sizes and the study results were drawn from multi-site data which are often heterogenous and used now outdated traditional low-resolution data acquisition protocols. In addition, there have been no human brain MRI studies that explore multiple psychiatric and neurological disorders that occur through the lifespan within the same cohort of subjects.

### 1.2. High-quality multi-modal MRI protocols and preprocessing pipelines

The HCP developed a broad approach to improving brain imaging data acquisition, preprocessing, analysis, and sharing (Glasser et al., 2016b). It includes: 1) high-quality multi-modal data acquisition; 2) in a large number of subjects; and 3) high-quality data preprocessing and has proven usefulness of MRI techniques for understanding the detailed organization of a healthy human brain (Elliott et al., 2018a; Glasser et al., 2016a; Smith et al., 2015). The HCP aimed to delineate the brain areas and characterize neural pathways that underlie brain function and behavior in 1,200 healthy young adults (Van Essen et al., 2012). HCP scans were performed by a single MR scanner (a customized 3T Skyra, Siemens Healthcare GmbH, Erlangen, Germany) in a total of 4-hour scan time for high-resolution multi-modal data, which included T1-weighted (T1w) images, T2-weighted (T2w) images, diffusion-weighted images (DWI), rsfMRI, and task fMRI (Glasser et al., 2016b; Glasser et al., 2013). The HCP also developed a set of preprocessing pipelines with improved cross-subject alignment that dramatically improves the spatial localization of brain imaging findings and also increasing statistical sensitivity (Coalson et al., 2018; Glasser et al., 2013; Robinson et al., 2018). For the Lifespan Developing and Aging HCP Projects (HCP-D and HCP-A) the original HCP protocol for healthy young adults was shortened, for children and the elderly (60 to 90 min scan time; (Bookheimer et al., 2019; Harms et al., 2018; Somerville et al., 2018), and for psychiatric and neurological disorders (the Connectomes Related to Human Disease [CRHD]; https://www.humanconnectome.org/disease-studies), and adolescent development (the ABCD project; (Casey et al., 2018). The UK Biobank used an even more abbreviated scanning approach to collect a much larger number of cohort (n = 100,000) to predict health conditions (Miller et al., 2016).

Many of these high-quality multimodal projects have been based on a single or small number of the same model scanners at different sites and thus did not fully address standardization of the data acquisition across different scanner models or vendors. We aim to accelerate harmonization technologies to be used in at least five scanner platforms by combining approaches to high-quality imaging acquisition, preprocessing, study design, and statistical bias correction to potentially improve the sensitivity and validity of imaging biomarkers.

### 1.3. Traveling subjects

A harmonization approach is required for individual-based statistics using a multi-site dataset, even when the brain images are obtained using the same machine and protocol, because the data from each site has the bias from hardware and scanning protocol (measurement bias) and sampling variability (i.e. age, sex, handedness, and socioeconomic status). If measurement biases were correlated or anti-correlated with a specific disease state this would result in a positive or negative bias in a given measure, whereas uncorrelated biases would merely reduce sensitivity (i.e. SNR) of the measure. Sampling biases due to biological differences in the sampled populations should also be considered for both case and control groups. Data harmonization has been proposed to control for these biases, including a general linear model (GLM) with the site as the covariate, a Bayesian approach (Fortin et al., 2018; Fortin et al., 2017), and a meta-analytic approach (Okada et al., 2016; van Erp et al., 2016), but the methods used for controlling both biases are unable to distinguish between them (Yamashita et al., 2019). Inter-site cross-validation by machine learning and deep learning techniques is a method that aims to remove bias without any specific preparation if large-sample datasets are available (Nunes et al., 2018). However, this method extracts stable characteristics across the images and is limited to using only a part of the information for further analysis. In addition, it is unclear whether the classifiers obtained by such methods can be applied to an independent new site of the initial multi-site project.

The traveling subject (TS) approach is a powerful research design to control for site differences (Figure 1). This approach requires the images from the same participants at all the participating sites, but also requires significant effort from the sites and the participants when compared to other harmonization methods listed above, and the TS scans must be completed before the analysis starts. However, the TS approach can differentiate most of the sample variability from measurement bias in functional MRI (Yamashita et al., 2019), structure and diffusion MRI (Tong et al., 2020). The DecNef Project explored rsfMRI functional connectivity for multiple psychiatric diseases and scanned nine TS participants who received repeated MRI measurements at all sites. Measurement and sampling biases for each group (schizophrenia, MDD, ASD, and healthy controls) were segregated from individual and disease-specific factors as the rest of sampling variability. The results showed that the effects of both bias types on functional connectivity were greater than or equal to those of disease-specific factors. With regard to measurement bias, differences in phase encoding direction had the biggest effect size when compared to those of vendor, coil, and scanner within the same vendor. The harmonization method was estimated to reduce measurement bias by 29% and improve the signal-to-noise ratio by 40% (Yamashita et al., 2019). Further investigations are needed to determine the best approach for reducing sampling bias arising from biological differences in the sampled population.

**Figure 1.**
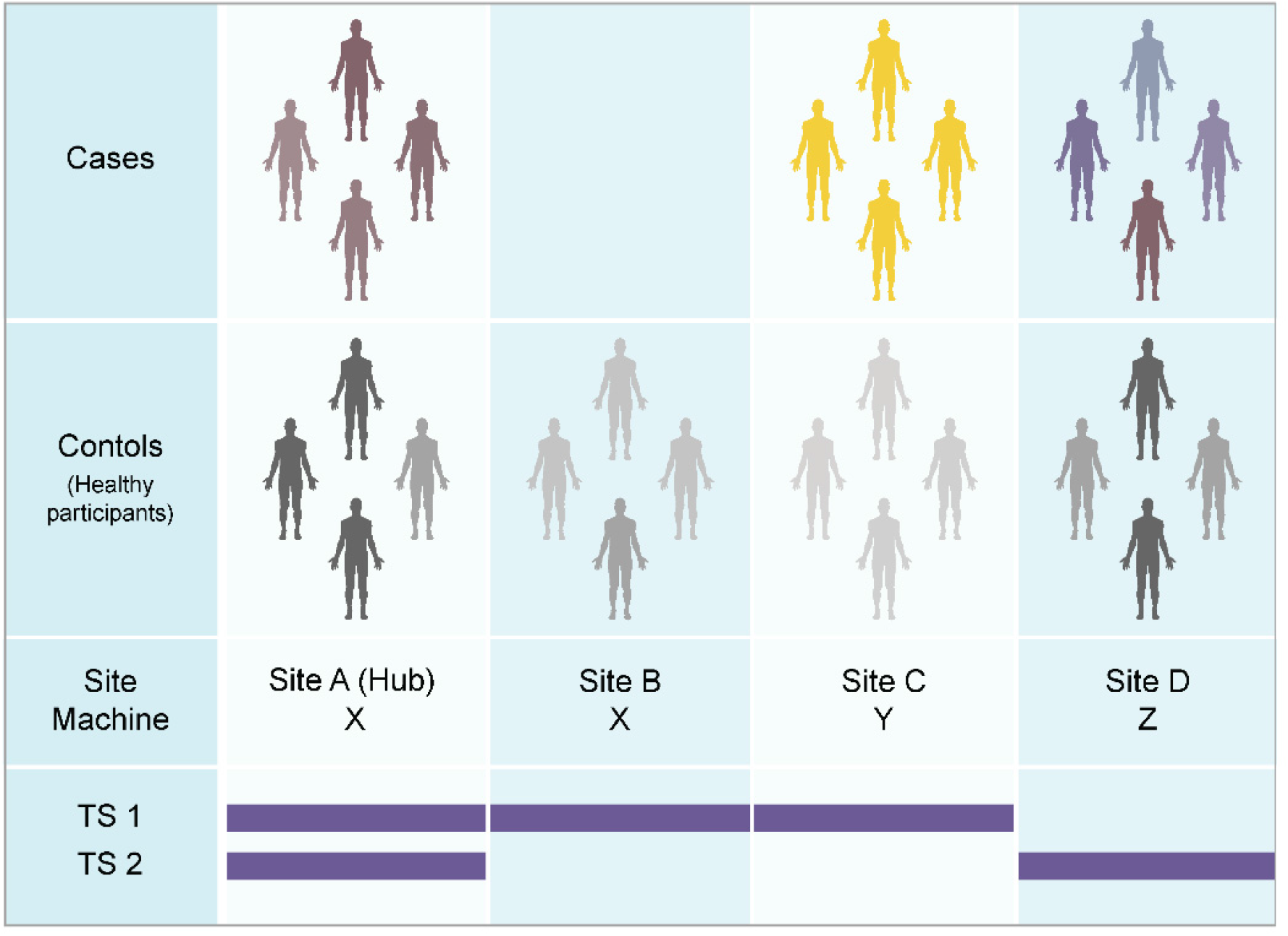
Case-control studies and traveling subject approach. (Top) When we analyze multi-site data from a set of case-control MRI studies, we must consider machine and protocol-derived bias (measurement bias) as well as sampling bias (from biological differences in the sampled populations). Even if the machine and protocol are the same between sites (e.g. Sites A and B), measurement bias may still occur because of slight differences in the magnetic or radiofrequency fields, etc. Sampling bias should be considered for patient groups as well as control groups, given that the control participants were recruited according to the demographics in the patient group. (Bottom) The traveling subject (TS) harmonization approach enables us to combine with case-control datasets by differentiating between measurement and sampling biases (Yamashita et al., 2019). Based on the general linear model (GLM), TS participants need to receive measurements from all participating sites (e.g. only TS 1 dataset). To reduce the effort of TS participants and participating sites, this project applies a general linear mixed model (GLMM) approach and hub-and-spoke model to the TS project. With this approach, all participants receive scans at one or more hub sites (site A), and measurement bias is calculated using multiple TS datasets by means of a GLMM (TS 1 and 2).

### 1.4. Brain/MINDS Beyond project

The Strategic International Brain Science Research Promotion Program (Brain/MINDS Beyond; FY2018–FY2023; https://brainminds-beyond.jp/) was funded by the Japan Agency for Medical Research and Development (AMED) to support global brain research by enhancing collaboration with the domestic projects of other countries. Brain/MINDS Beyond consists of four research groups: G1-1, Identification of the pathogenic mechanism of psychiatric and neurological disorders through the acquisition and analysis of brain MRI-scan images and clinical data (Developmental [G1-1D], adult [G1-1A], and senescent [G1-1S] stages); G1-2, Brain MRI data acquisition, analysis, and informatics; G2, Research involving an inter-species comparison of human and nonhuman primate brains by structural and functional parcellation and homology analyses; and G3, Development and application of technologies, such as neuro-feedback through collaboration with artificial intelligence research projects as well as the Innovative Research Group. In human brain imaging, G1-1 intends to measure human participants, including patients with neuropsychiatric disorders, across the lifespan, and G1-2 intends to coordinate and support data acquisition, storage, preprocessing, analysis, and distribution (Figure 2 and Table 1). The Brain/MINDS Beyond MRI working group also set up a standardized procedure for MRI data acquisition (Harmonization protocol [HARP]) and clinical and neurocognitive data assessment (Tables 2 and 3). Following previous multi-site studies in Japan (Iwatsubo et al., 2018; Okada et al., 2016; Yahata et al., 2016; Yamashita et al., 2019), the overall goal of this project is expected to find altered brain imaging characteristics in psychiatric and neurological disorders that can be applied to future therapeutic investigations and clinical devices. To address limitations of previous findings in multi-site studies, we are using high performance research-based MRI scanners and we modeled our multi-modal protocol (T1w images, T2w images, diffusion-weighted imaging [DWI], rsfMRI, task fMRI, quantitative susceptibility mapping, and arterial spin labeling) on that used by the HCP and ABCD study projects. We are also obtaining a TS dataset for the harmonization of the clinical MRI datasets and the development of technical tools to harmonize the multi-site data. Once the project period ends, the data will be openly distributed to researchers via a public database.

**Table 1.**
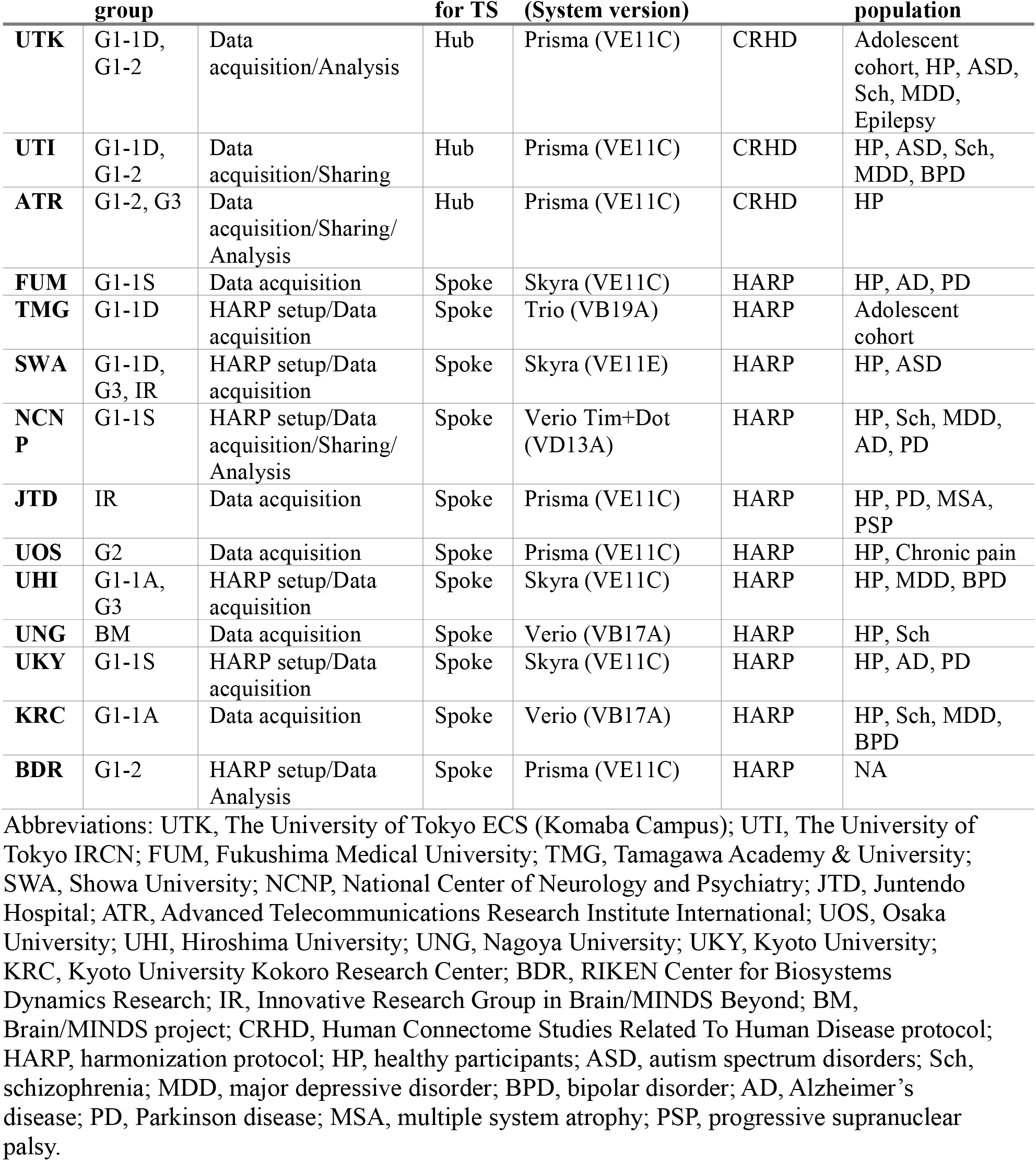
Participating sites of the Brain/MINDS Beyond MRI project.

| Site | Research group | Role for the project | Role for TS | MRI scanner (System version) | Protocol | Main target population |
| --- | --- | --- | --- | --- | --- | --- |
| UTK | G1-1D, G1-2 | Data acquisition/Analysis | Hub | Prisma (VE11C) | CRHD | Adolescent cohort, HP, ASD, Sch, MDD, Epilepsy |
| UTI | G1-1D, G1-2 | Data acquisition/Sharing | Hub | Prisma (VE11C) | CRHD | HP, ASD, Sch, MDD, BPD |
| ATR | G1-2, G3 | Data acquisition/Sharing/Analysis | Hub | Prisma (VE11C) | CRHD | HP |
| FUM | G1-1S | Data acquisition | Spoke | Skyra (VE11C) | HARP | HP, AD, PD |
| TMG | G1-1D | HARP setup/Data acquisition | Spoke | Trio (VB19A) | HARP | Adolescent cohort |
| SWA | G1-1D, G3, IR | HARP setup/Data acquisition | Spoke | Skyra (VE11E) | HARP | HP, ASD |
| NCNP | G1-1S | HARP setup/Data acquisition/Sharing/Analysis | Spoke | Verio Tim+Dot (VD13A) | HARP | HP, Sch, MDD, AD, PD |
| JTD | IR | Data acquisition | Spoke | Prisma (VE11C) | HARP | HP, PD, MSA, PSP |
| UOS | G2 | Data acquisition | Spoke | Prisma (VE11C) | HARP | HP, Chronic pain |
| UHI | G1-1A, G3 | HARP setup/Data acquisition | Spoke | Skyra (VE11C) | HARP | HP, MDD, BPD |
| UNG | BM | Data acquisition | Spoke | Verio (VB17A) | HARP | HP, Sch |
| UKY | G1-1S | HARP setup/Data acquisition | Spoke | Skyra (VE11C) | HARP | HP, AD, PD |
| KRC | G1-1A | Data acquisition | Spoke | Verio (VB17A) | HARP | HP, Sch, MDD, BPD |
| BDR | G1-2 | HARP setup/Data Analysis | Spoke | Prisma (VE11C) | HARP | NA |

Abbreviations: UTK, The University of Tokyo ECS (Komaba Campus); UTI, The University of Tokyo IRCN; FUM, Fukushima Medical University; TMG, Tamagawa Academy & University; SWA, Showa University; NCNP, National Center of Neurology and Psychiatry; JTD, Juntendo Hospital; ATR, Advanced Telecommunications Research Institute International; UOS, Osaka University; UHI, Hiroshima University; UNG, Nagoya University; UKY, Kyoto University; KRC, Kyoto University Kokoro Research Center; BDR, RIKEN Center for Biosystems Dynamics Research; IR, Innovative Research Group in Brain/MINDS Beyond; BM, Brain/MINDS project; CRHD, Human Connectome Studies Related To Human Disease protocol; HARP, harmonization protocol; HP, healthy participants; ASD, autism spectrum disorders; Sch, schizophrenia; MDD, major depressive disorder; BPD, bipolar disorder; AD, Alzheimer's disease; PD, Parkinson disease; MSA, multiple system atrophy; PSP, progressive supranuclear palsy.

**Table 2.**
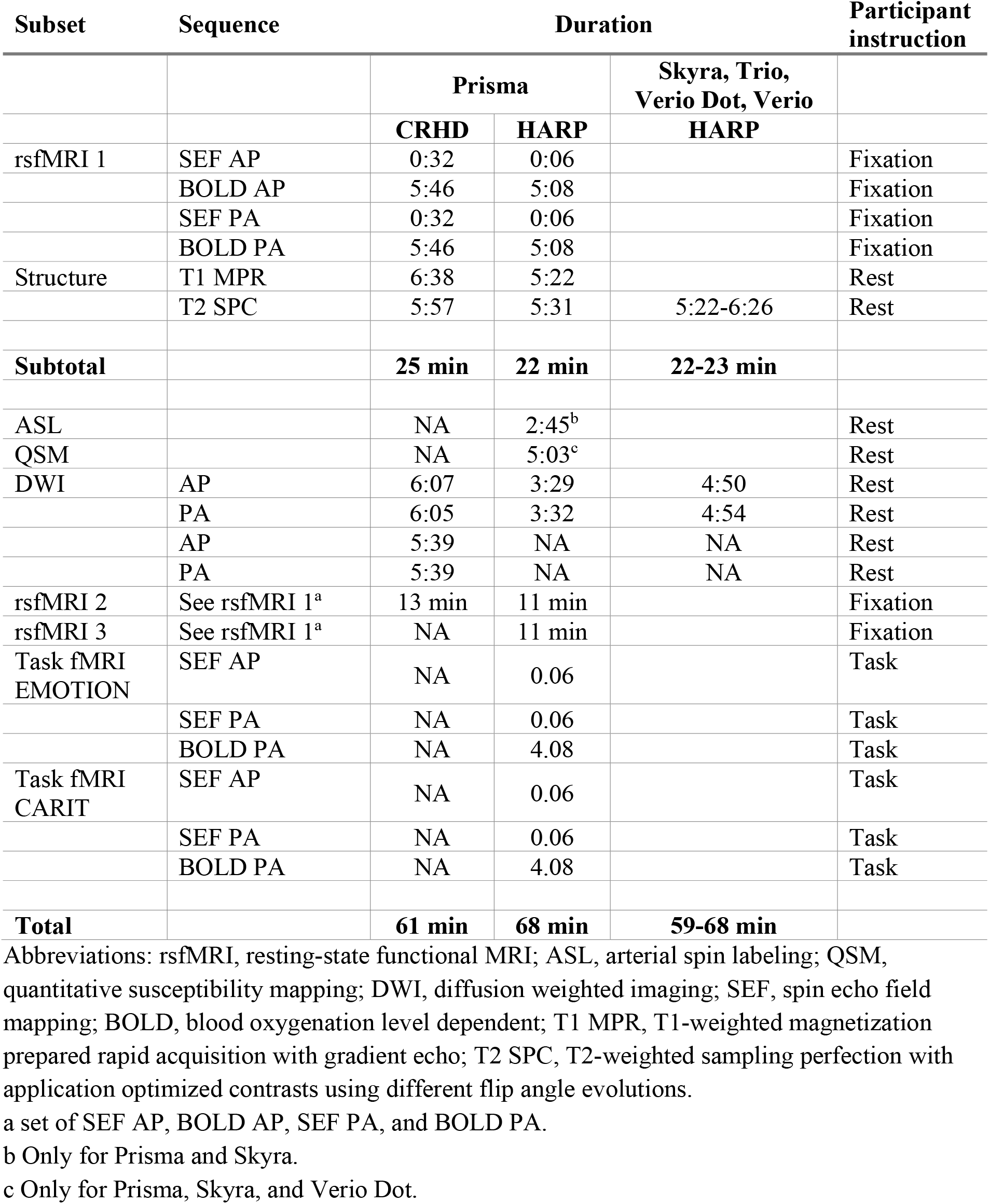
CRHD and HARP protocols.

| Subset | Sequence | Duration |  |  | Participant instruction |
| --- | --- | --- | --- | --- | --- |
|  |  | Prisma |  | Skyra, Trio, Verio Dot, Verio |  |
|  |  | CRHD | HARP | HARP |  |
| rsfMRI 1 | SEF AP | 0:32 | 0:06 |  | Fixation |
|  | BOLD AP | 5:46 | 5:08 |  | Fixation |
|  | SEF PA | 0:32 | 0:06 |  | Fixation |
|  | BOLD PA | 5:46 | 5:08 |  | Fixation |
| Structure | T1 MPR | 6:38 | 5:22 |  | Rest |
|  | T2 SPC | 5:57 | 5:31 | 5:22-6:26 | Rest |
| Subtotal |  | 25 min | 22 min | 22-23 min |  |
| ASL |  | NA | 2:45<sup>b</sup> |  | Rest |
| QSM |  | NA | 5:03<sup>c</sup> |  | Rest |
| DWI | AP | 6:07 | 3:29 | 4:50 | Rest |
|  | PA | 6:05 | 3:32 | 4:54 | Rest |
|  | AP | 5:39 | NA | NA | Rest |
|  | PA | 5:39 | NA | NA | Rest |
| rsfMRI 2 | See rsfMRI 1<sup>a</sup> | 13 min | 11 min |  | Fixation |
| rsfMRI 3 | See rsfMRI 1<sup>a</sup> | NA | 11 min |  | Fixation |
| Task fMRI EMOTION | SEF AP | NA | 0.06 |  | Task |
|  | SEF PA | NA | 0.06 |  | Task |
|  | BOLD PA | NA | 4.08 |  | Task |
| Task fMRI CARIT | SEF AP | NA | 0.06 |  | Task |
|  | SEF PA | NA | 0.06 |  | Task |
|  | BOLD PA | NA | 4.08 |  | Task |
| Total |  | 61 min | 68 min | 59-68 min |  |

Abbreviations: rsfMRI, resting-state functional MRI; ASL, arterial spin labeling; QSM, quantitative susceptibility mapping; DWI, diffusion weighted imaging; SEF, spin echo field mapping; BOLD, blood oxygenation level dependent; T1 MPR, T1-weighted magnetization prepared rapid acquisition with gradient echo; T2 SPC, T2-weighted sampling perfection with application optimized contrasts using different flip angle evolutions.
<sup>a</sup> a set of SEF AP, BOLD AP, SEF PA, and BOLD PA.
<sup>b</sup> Only for Prisma and Skyra.
<sup>c</sup> Only for Prisma, Skyra, and Verio Dot.

**Table 3.** Clinical and neuropsychological assessment.

|  | G1-1D | G1-1A | G1-1S |
| --- | --- | --- | --- |
| Depression | K6 or BDI-II | BDI-II and PHQ-9 | PHQ-9 and BDI-II/GDS-15 |
| Anxiety | — | GAD-7 | STAI |
| Autism | AQ-10, AQ-50 or SRS-2 (for developmental disorders) | AQ-10 or AQ-50 | — |
| Psychosis | APSS | — | NPI-Q |
| Intellectual ability | JART-25 or WAIS-III (WISC at the age of 15 years or less) Information and Picture completion subtests | JART-25 | JART-25 |
| Cognitive function | CANTAB or BACS-J | CANTAB or BACS-J | ADAS-Cog11, CDT, CDR, FAB, HVLT-R, JLO, MMSE, MoCA-J, SDMT, TMT-A/B, WMS-R |
| General function and disability | GAF, mGAF or WHO-DAS 2.0 | GAF, mGAF or WHO-DAS 2.0 | Schwab & England ADL |
| Quality of life | EQ-5D | EQ-5D | PASE |
| Well-being | WHO-5 | WHO-5 | SHAPS |
| Handedness | EHRS or UTokyo | EHRS or UTokyo | UTokyo |

**Figure 2.**
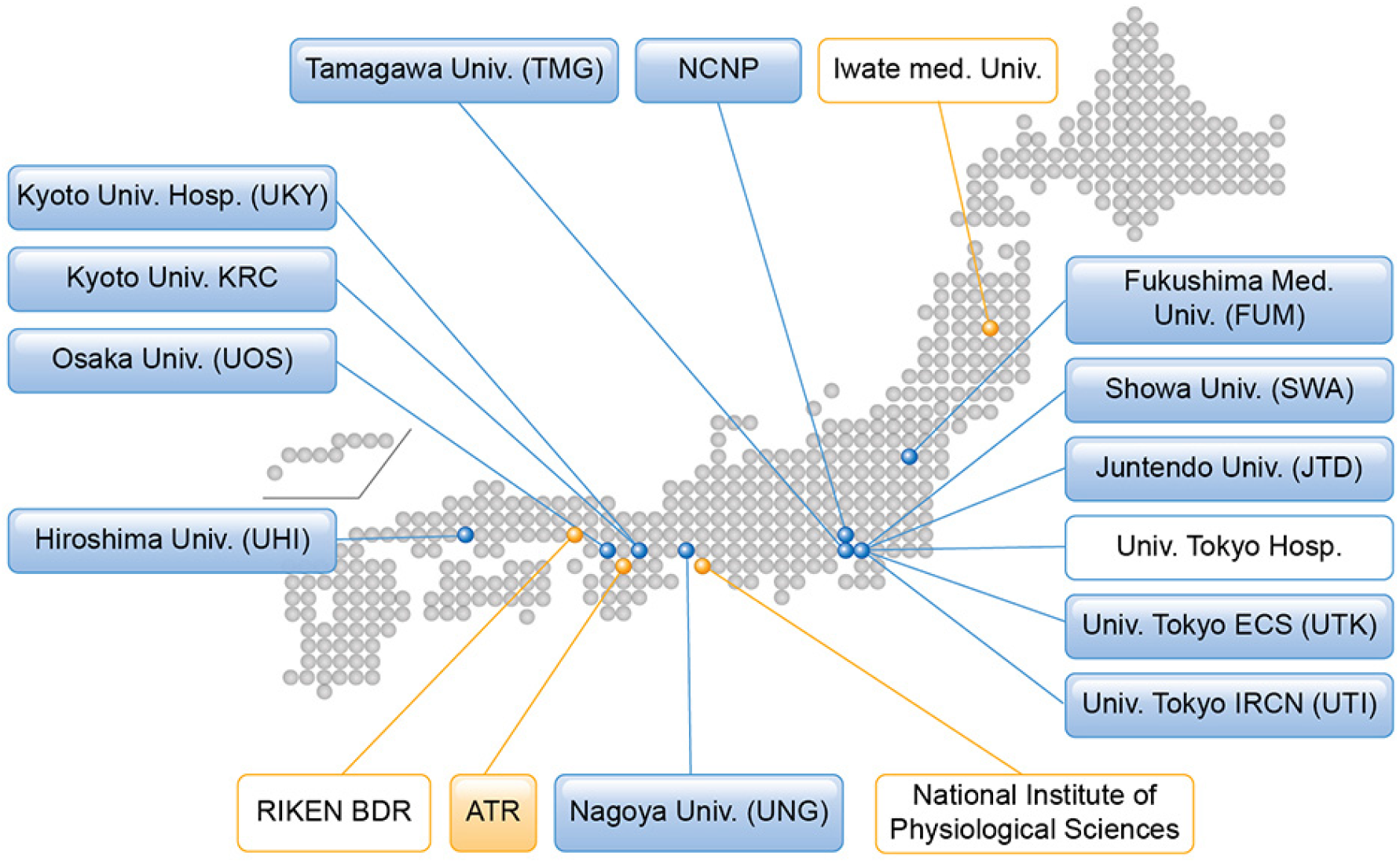
Brain/MINDS Beyond human brain MRI project. Institutes in the blue boxes show measurement and analysis sites for neuropsychiatric disorders, and those in the orange boxes show analysis support sites. Institutes listed in boxes with a colored background represent participation in the traveling subject project.

Here, we introduce the Brain/MINDS Beyond human brain MRI project and show preliminary results in high-quality neuroimaging using the TS data that is amenable to harmonization. We then discuss our plans for investigating the neural basis of psychiatric and neurological disorders in the hope of developing therapeutic targets and devices that are applicable to clinical settings.

## 2. Brain/MINDS Beyond human brain MRI study

### 2.1. Participating sites and target population

As of March 2020, 13 sites have approved this study project, received approval from their respective ethical review board(s), and obtained clinical and TS measurements using the appropriate MRI scanners (Table 1). Of these, 5 sites mainly explore psychiatric disorders (schizophrenia, ASD, MDD, and BPD), 4 sites neurological disorders (AD, PD, multiple system atrophy, progressive supranuclear palsy, chronic pain disorder, and epilepsy), and 2 sites both categories. Two sites measure the general adolescent population to investigate brain development and recruit through advertisement and cohort studies (Ando et al., 2019; Okada et al., 2019). Each site intends to obtain brain images and demographic (and clinical) characteristics for clinical cases and match controls for age, sex, premorbid IQ or educational attainment, socio-economic status, and handedness (See Cognitive and behavioral assessment section). The exclusion criteria were set by each study purpose (i.e. low premorbid IQ, history of loss of consciousness for more than 5 min, illegal drug use, and alcohol dependency). Illegal drug use can be a major concern for disease onset and poor prognosis, especially for psychiatric disorders. However, there is far less illegal drug use in Japan compared to Western European countries (Degenhardt et al., 2008; Lee and Kwon, 2016), and most of the participating sites excluded those with a current illegal drug use or previous history of regular use (Koike et al., 2013).

For the TS project, 75 healthy adults planned to undergo 6 to 8 scans at three or more sites within 6 months (See Traveling Subject Project section). Five or more participants per site were recruited. Each participant received test-retest scans at the recruitment site and underwent scans at different sites including a hub site. We set up three hub sites, according to a hub-and- spoke model, in which all participants received scans using a MAGNETOM Prisma scanner (Siemens Healthcare GmbH, Erlangen, Germany) and the CRHD and HARP protocols.

### 2.2. Harmonized brain MRI protocols

We developed protocols that minimize potential differences related to measurement and increase the MR image sensitivity to brain organization in psychiatric and neurological disorders. From a neurobiological perspective, the cerebral cortex is organized by a 2D sheet-like structure with an average thickness of 2.6 mm embedded and folded in the ∼1300 mL of brain volume (Glasser et al., 2016b). From a neuroimaging perspective, the spatial resolution and homogeneity of the images are important factors that may induce bias and error during the image analysis; these include partial voluming, image distortion, errors in brain segmentation, and registration. Of these, respecting spatial fidelity of neuroanatomical structures is the most important approach for achieving unbiased imaging (Glasser et al., 2016b). Therefore, the spatial resolution of the imaging was determined based on cortical thickness and was matched across all scanners. The phase encoding direction of EPI-based functional and diffusion MRI is an important factor that relates to spatial distortion (and signal loss in fMRI) in association with the polarity of the direction, echo spacing, and B0 magnetic field homogeneity; therefore, we acquire a spin-echo filed map with opposite phase encoding directions to enable distortion correction (Andersson et al., 2003). Based on these strategies, two MRI protocols were planned for use in the project: 1) a harmonized MRI protocol (HARP), which can be run on the multiple MRI scanners/sites within a period of 22 to 65 min; and 2) an ‘HCP style’ MRI protocol used by HCP CRHD for the high-performance 3T MRI scanner (e.g. MAGNETOM Prisma).

The HARP was created to be used at multiple MRI scanners/sites, and it was designed to obtain high-quality and standardized brain MRI data in a ‘clinically’ practical window of time (Table 2 and Supplementary Table S1). The parameters of the MRI scanners were as follows: 1) static magnetic field strength of 3T; 2) multi-array head coil with 32 or more channels; and 3) ability to perform a multi-band EPI sequence provided from Center for Magnetic Resonance Research, University of Minnesota with an acceleration factor of 6 (Moeller et al., 2010; Setsompop et al., 2012; Xu et al., 2013). In 2019, the protocol was adapted for use with five MRI scanners/systems (MAGNETOM Prisma, Skyra, Trio A Tim, Verio, and Verio Dot; Siemens Healthcare GmbH, Erlangen, Germany), and we plan to expand it to different MRI scanners/vendors during the project period and in fact we are working on creating HARP protocol for GE scanners. The HARP was intended to perform the brain scan within a period of ∼ 30 min using a high-resolution structural MRI scan (T1w and T2w, spatial resolution of 0.8 mm) and two high-sensitive rsfMRI scans with opposing phase directions, a spatial resolution of 2.4 mm, and a temporal resolution of 0.8 s for a total of 10 minutes. The protocols also include optional sequences for four additional rsfMRI scans, task fMRI (Emotion and CARIT) (Winter and Sheridan, 2014), two DWI scans with opposing phase encoding directions, quantitative susceptibility mapping, and arterial spin labeling. The minimum and maximum scanning time of the HARP is 22 and 65 min, respectively (Table 2). The preliminary results across scanners and multi-array coils in the same subject (ID = 9503) revealed that the temporal signal-to-noise ratio (tSNR) was very high in all the scanners. The mean ± standard deviation across 32k greyordinates was 161 ± 80 in the Prisma at UTK, 155 ± 81 in the Verio Dot at SWA, 151 ± 72 in the Skyra fit at SWA, 151 ± 80 in the Verio at ATR, and 150 ± 74 in the Prisma fit at ATR; the values and their distributions were similar across scanners/sites (Figure 3A).

**Figure 3.**
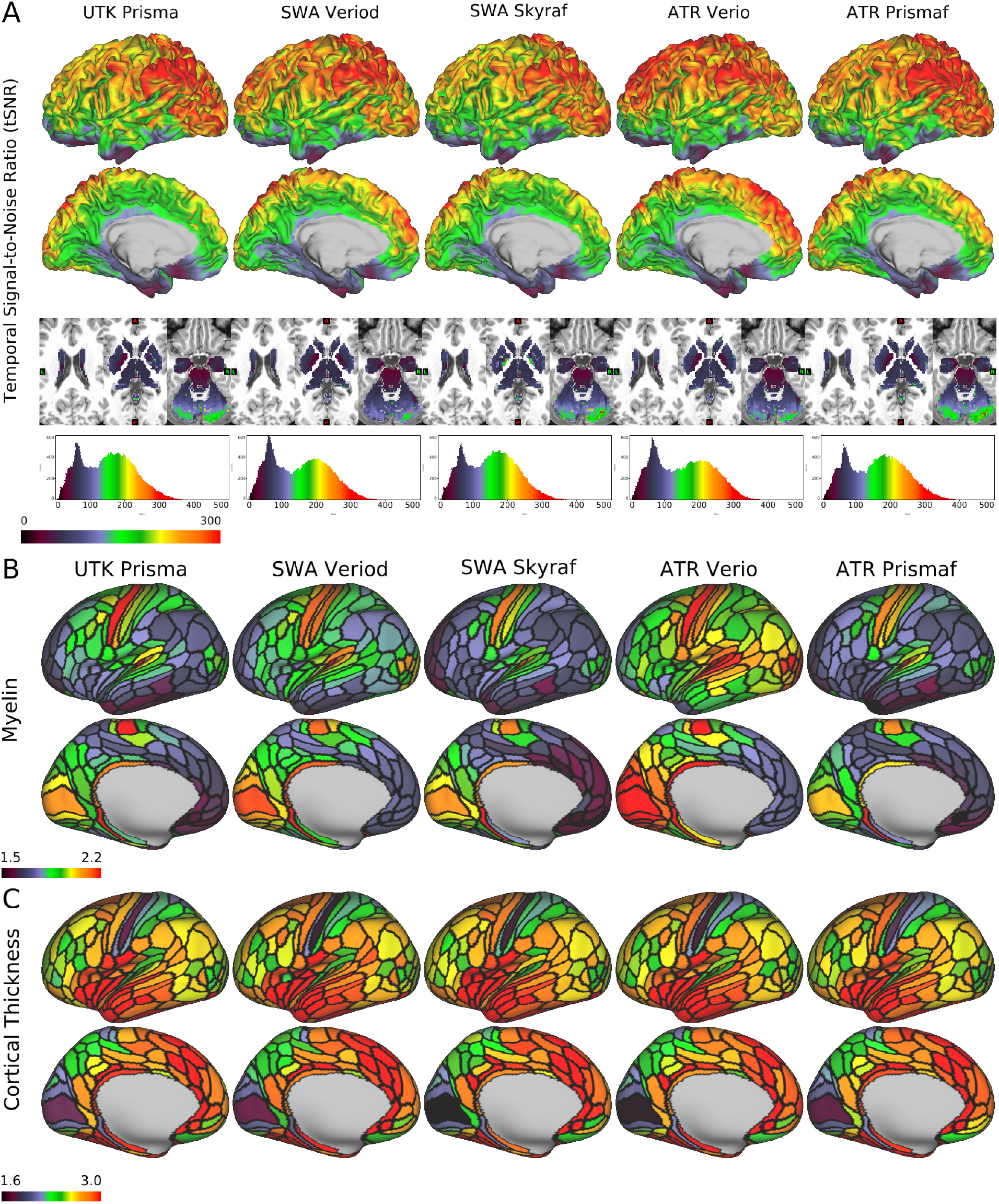
Quality of MRI and preliminary cortical structures obtained by HARP in a single traveling subject across scanners/sites. A) Temporal signal-to-noise ratio (tSNR) obtained in a single subject (ID = 9503) across different scanners/sites by a harmonized MRI protocol (a sequence of functional MRI in HARP using a multi-band echo planar imaging with TR/TE = 800/34.4 ms; see Supplementary Table S1 for other details). The images from top to bottom show color-coded tSNR maps in 32k greyordinates (see main text) overlaid on the lateral and medial surface of the mid-thickness surface of the left hemisphere, the subcortical sections of the T1w image, and the histogram of the tSNR values. B) Cortical myelin contrast (T1w/T2w ratio) across different scanners. The myelin contrast is not corrected for the biasfield and parcellated by the HCP MMP v1.0 (Glasser et al., 2016a). C) The map shows cortical thickness across different scanners. Cortical thickness is corrected by curvature and parcellated by the HCP MMP v1.0. The tSNR, myelin map and cortical thickness are comparable across scanners. Data at https://balsa.wustl.edu/7q4P9 and https://balsa.wustl.edu/6Vvqv

The CRHD protocol was planned for collaboration with the HCP CRHD for the Early Psychosis Project. The HCP CRHD protocol also included high-resolution structural MRI (spatial resolution of 0.8 mm), high-resolution resting-state fMRI (spatial resolution of 2 mm) with an opposing phase encoding direction and longer scan time, and high-resolution and high angular diffusion MRI.

The installation of the protocols in the MRI scanners was ensured by conducting hierarchical parameter checks and site visits at the beginning of the measurement period. After the protocol installation, each site sent XML files of the installed protocol from the MRI scanner to the protocol management site (UTK), and all the parameters were confirmed with a checksum algorithm using R (R Core Team, 2018). This process was useful for validating the protocols across sites/scanners because some of the MRI scanners actually underwent inappropriate installation and were set with different parameters. The results were then sent back to the collaborators, who edited the parameters. We also checked the DICOM files that are deposited in the ATR XNAT server. In this phase, we checked the parameters, slice numbers, and diffusion gradient information (bvec and bval).

The manuals were shared and used at the sites for protocol installation, demographic and clinical assessment before the scan (e.g. handedness), and the assessment of and instruction to participants during the scan (e.g. general instruction during the scan, fixation to the cross during rsfMRI scans, and the assessment of sleepiness during the rsfMRI).

### 2.3. Cognitive and behavioral assessment

Each participating site assesses demographic characteristics (i.e. age, sex, and socioeconomic status), clinical characteristics (i.e. diagnosis, symptom severity, cognitive function, and general functioning), and subjective social evaluations (i.e. quality of life and well-being) (Table 3). Each subgroup (G1-1D, G1-1A, G1-1S, and G1-2 TS) indicates standard scales, some of which are uniform across subgroups and easier to share and use when analyzing brain images.

### 2.4. Travelling subject project

Based on the previous study (Yamashita et al., 2019), we conduct a TS project for the CRHD and HARP protocols. At some sites, we also scan TS with the previous protocol (named as SRPB [Strategic Research Program for Brain science]), which was used in the multi-site studies to achieve retrospective harmonization (Iwatsubo et al., 2018; Okada et al., 2016; Yahata et al., 2016; Yamashita et al., 2019). Because we limit the scanners, head coils, and protocols in this project, we expect to see reduced measurement biases, which may enhance the disease-related effect size in clinical studies and provide better ways to diminish bias in future studies.

The previous data harmonization using the TS dataset was based on a GLM (Yamashita et al., 2019), in which participants needed to travel to all the sites/scanners. In contrast, the present TS project was designed so that the participants travel only some of the test sites/scanners, and statistical harmonization flexibly adapt the incompleteness by using a general linear mixed model (GLMM; Figure 1). This design may also adapt the incompleteness of the scanning for each participant, since the current protocols require a longer scan time compared to previous ones, and thus result in potential cancellation or data completeness. To ensure harmonization across all the sites/scanners, we applied a hub-and-spoke model to arrange traveling scans at each recruitment site (Supplementary Table S2). Each participant undergoes CRHD and HARP scans using the Prisma (∼2 hours) at one or more hub sites (UTK, UTI, and ATR) to harmonize the data within the Brain/MINDS Beyond project and other projects (e.g. Brain/MINDS, HCP, and ABCD) and test the difference in quality between the protocols. The other visiting sites were determined in consideration of the site locations, machine differences, and project similarities between the sites. Each participant receives multiple scans at the recruitment site to assess the test-retest reliability (1-hour x 2 sessions).

For the TS project, 75 healthy adults—five or more participants per site—are scheduled to undergo 6 to 8 scans at three or more sites within 6 months (Supplementary Table S2). The total number of scans and spokes between the sites are expected to be 455 and 465, respectively (Figure 4A). As of March 2020, 74 participants were registered and 405 scans (89.0 %) were completed and uploaded to the ATR XNAT server. The data provided 368 spokes (76.1 %, Figure 4B). The TS project will end in August 2020.

**Figure 4.**
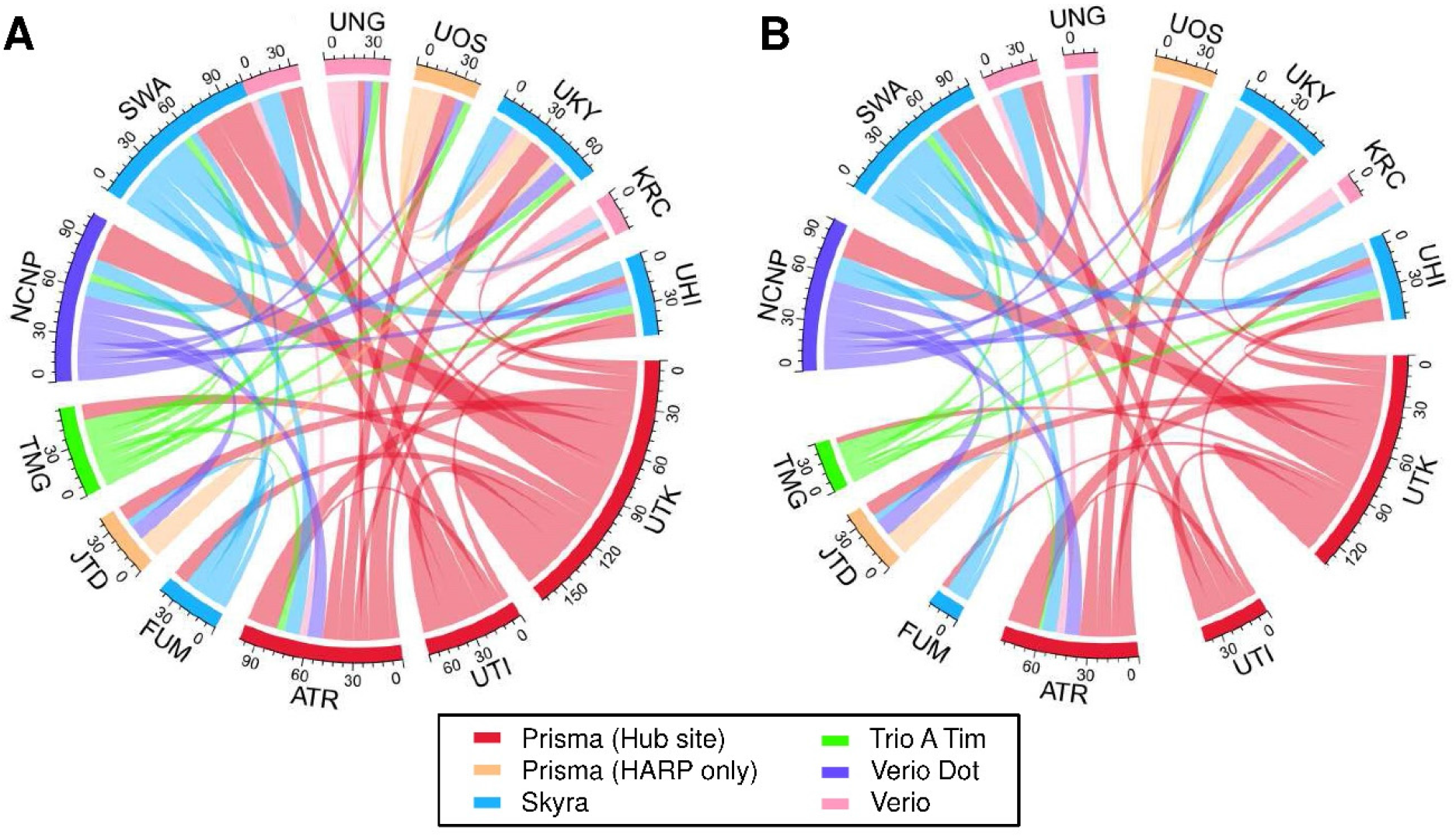
Expected and current data connection of the traveling subjects. Data connections in the traveling subject project that were (A) initially planned and (B) the actual connections as of March 2020. Hub sites using Prisma and other sites using Prisma, Skyra, Trio A Tim, Verio Dot, and Verio are illustrated in red, orange, blue, green, purple, and pink, respectively.

### 2.5. Data storage, preprocessing, and quality control

#### 2.5.1. Data logistics

Brain MR images obtained using the CRHD and HARP protocols in this study project and related studies are stored, preprocessed, and distributed using the XNAT server system (https://www.xnat.org/) (Figure 5). Due to the legacy of previous multi-site studies (Iwatsubo et al., 2018; Yahata et al., 2016; Yamashita et al., 2019), several data centers were already available for this project. The images obtained from the development and adult projects (G1-1D and G1-1A) will be sent to an XNAT server at ATR and the clinical data will be sent to UTI. For the senescent project (G1-1S), all the data will be sent to the NCNP (Iwatsubo et al., 2018). The TS data will also be sent to the ATR server shown in dashed lines. When uploading to the XNAT server, personal information (i.e. name and date of birth) contained in DICOM is automatically removed using an anonymization script of XNAT. A defacing procedure is performed for T1w and T2w images. These processes de-identify the MRI data. After manually checking whether the face images are completely obscured, all the anonymized MRI data are shared using Amazon AWS with RIKEN BDR, in which all image preprocessing is performed (See Preprocessing pipelines section). Preprocessed data are sent back to the servers and can be seen with limited access (i.e. participating sites). After a quality control (QC), cleaned imaging data with a demographic and clinical datasheet will be stored in the distribution server(s). All data will be also sent to backup server(s).

**Figure 5.**
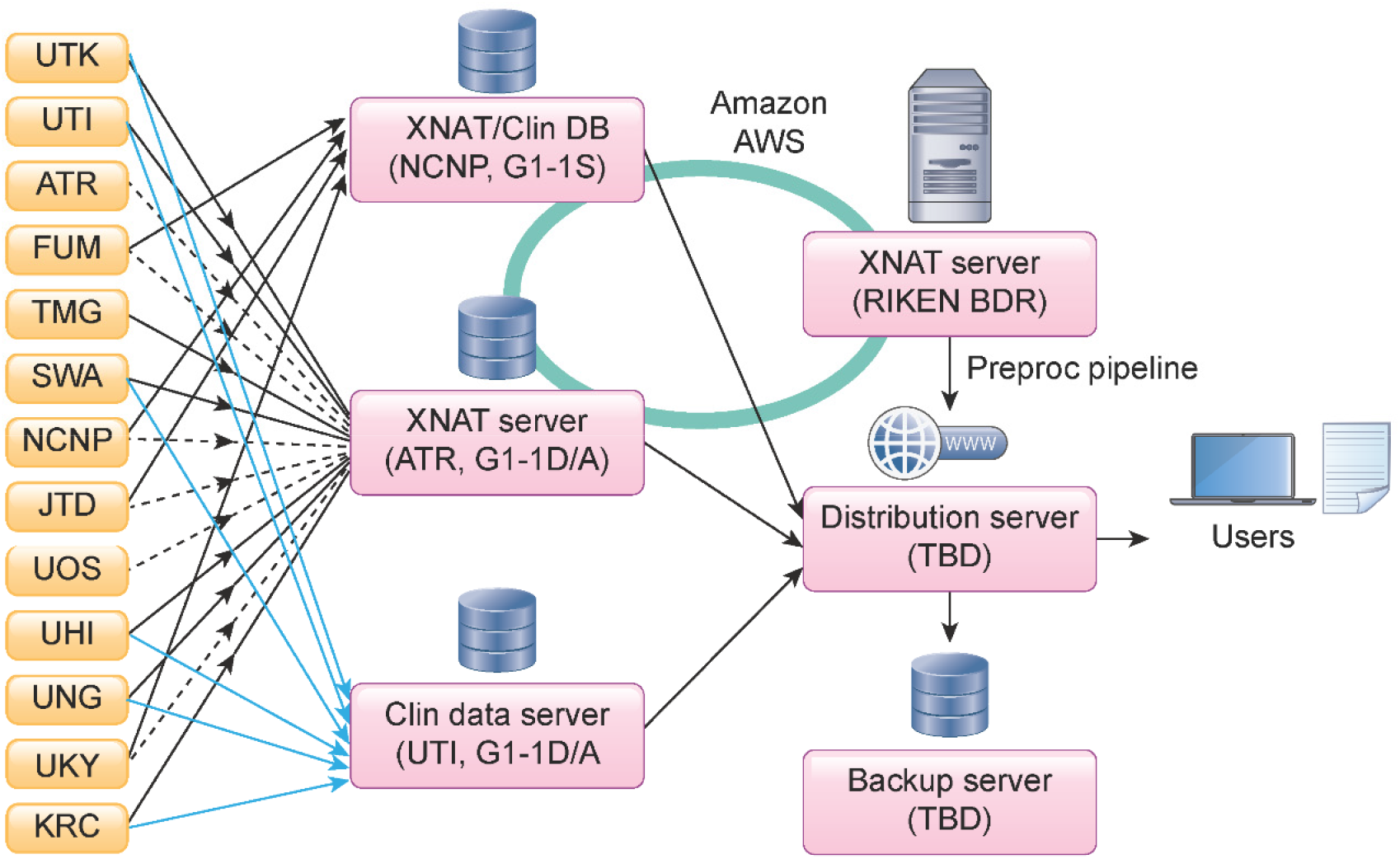
Data storage, preprocessing, quality check, and data sharing. MRI (black line) and clinical (blue line) data from G1-1D and G1-1A sites are sent to the XNAT server and a data server at ATR and UTI, respectively. All data from G1-1S sites are sent to an XNAT server and a data server managed by NCNP, as this group applied a standard clinical assessment protocol to the project following a previous multi-site study. Traveling subject data from G1-1S sites are also sent to the XNAT server in ATR (dot line). XNAT servers at NCNP, ATR, and RIKEN BDR are linked by Amazon AWS to share the imaging data. NCNP manages a separate server for storing clinical data (Clin DB) being collected from the participants in this project. All MR images are preprocessed at RIKEN BDR. All MR images are preprocessed at RIKEN BDR. All the raw and preprocessed data will be stored and provided to the users in a distribution server. A backup server will be placed at a different site.

#### 2.5.2. Preprocessing pipelines

All neuroimaging data are preprocessed at RIKEN BDR for this project. The MR images are sent via Amazon S3 to a high-throughput parallel computing system at RIKEN BDR for preprocessing. The raw MRI data in DICOM format are converted to those in NIFTI using a conversion program, BCILDCMCONVERT (https://github.com/RIKEN-BCIL/BCILDCMCONVERT), by which folder structures are created and all the imaging parameters are read and stored including the type of gradient, k-space read out time in phase and read directions, phase encoding directions, to be used for preprocessing. The preprocessing is performed using the HCP pipeline 4.2.0 (Glasser et al., 2013) with modifications for adapting and harmonizing multiple scanners. In brief, the structural MRI (T1w and T2w) is first corrected for image distortions related to the gradient nonlinearity in each scanner type and the inhomogeneity of the B0 static magnetic field in each scan. The signal homogeneity is dealt with by prescan normalization and is also improved by a biasfield correction using T1w and T2w images (Glasser and Van Essen, 2011). The T1w and T2w images are fed into non-linear registration to the Montreal Neurological Institute (MNI) space and used for cortical surface reconstruction using FreeSurfer (Fischl, 2012), surface registration using multi-modal surface matching (MSM) (Robinson et al., 2018) and folding pattern (MSMsulc); this is followed by the creation of a myelin map using T1w divided by T2w and surface mapping (Glasser and Van Essen, 2011). An example of a cortical myelin map (not biasfiled corrected [non BC]) in a single subject (ID = 9503) across scanners/sites is parcellated by HCP MMP v1.0 (Glasser et al., 2016a) and presented in Figure 3B, revealing the typical cortical distribution of the high myelin contrast in the primary sensorimotor (aeras 1, 3a, 3b, 4), auditory (A1), visual (V1), middle temporal, and ventral prefrontal (47m) areas—as demonstrated previously (Glasser and Van Essen, 2011). The distributions over the cortex were comparable between scanners, although absolute values were slightly different suggesting the residual bias from transmit field across scans/scanners (see also 2.5.3).

The functional MRI data is corrected for distortion (gradient nonlinearity and B0-inhomogeneity) and motion. The distortion from B0 static field inhomogeneity is corrected by means of opposite phase encoding spin echo fieldmap data using TOPUP (Andersson et al., 2003); it is then warped and resampled to MNI space at a 2 mm resolution and saved as a volume in the Neuroimaging Informatics Technology Initiative (NIFTI) format. The region of the cortical ribbon in the fMRI volume is further mapped onto the cortical surface and combined with voxels in the subcortical gray region to create 32k greyordinates in the Connectivity Informatics Technology Initiative (CIFTI) format. Multiple runs of the fMRI data are merged and fed into independent component analyses (ICA) followed by an automated classification of noise components and the removal of noise components using FIX (Salimi-Khorshidi et al., 2014) (Glasser et al., 2018). The automated classifier is trained using the data in this project and its accuracy is maximized. The denoised fMRI data, in combination with other cortical metrics (myelin, thickness; Figure 3B and 3C, respectively), is further used for multi-modal registrations (MSMAll) over the cortical surface, followed by ‘de-drifting’ (removing registration bias after multimodal registration) based on the group sampled in this study (Glasser et al., 2016a). The resting-state seed-based functional connectivity in an example of a single subject (ID = 9503) revealed a typical pattern over the cerebral cortex across scanners/sites; the left frontal eye field (FEF)-seed functional connectivity showed symmetric coactivation in the bilateral premotor eye field (PEF) (Figure 6A), whereas the left area 55b-seed FC showed an asymmetric language network distributed in the peri-sylvian language (PSL) area, superior temporal sulcus (STS), and areas 44/45 predominantly in the left hemisphere (Figure 6B).

**Figure 6.**
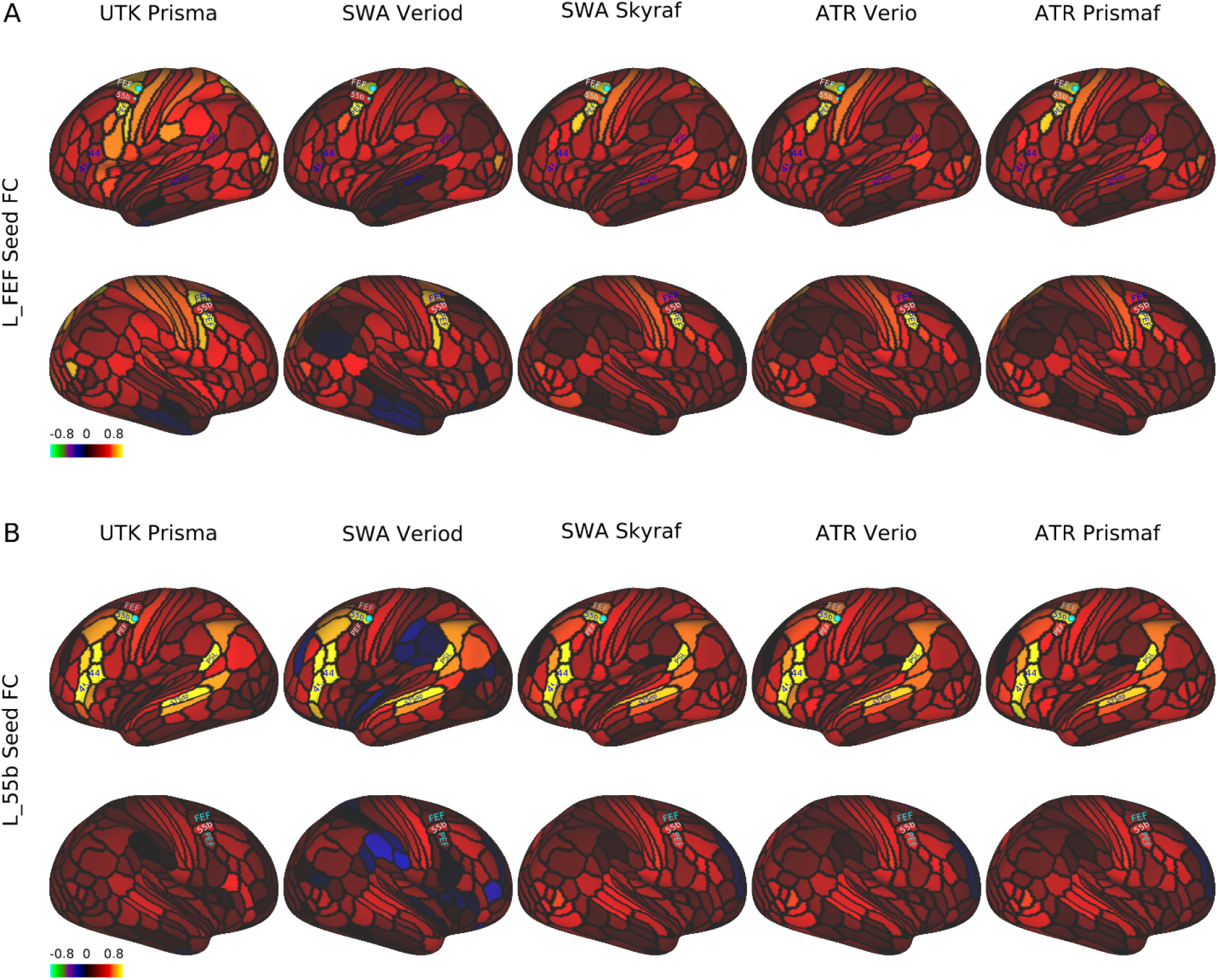
Seed-based resting-state functional connectivity in a single traveling subject across scanners/sites. In a single subject (ID = 9503), the resting-state fMRI scans (5 min x 4) were collected using a scanning protocol of HARP across different scanners/sites (see Supplementary Table S1), preprocessed, and denoised by a surface-based analysis to generate parcellated functional connectivity (FC) using the HCP MMP v1.0 (Glasser et al., 2016a). A) FC seeded from the left frontal eye field (FEF), which was distributed symmetrically in the bilateral premotor eye field (PEF) and comparable across scanners/sites. B) FC seeded from the left area 55b, which showed an asymmetric language network predominant in the left hemisphere that was comparable across scanners/sites. The language network is distributed in the areas of 44/45, superior temporal sulcus, dorsal posterior part (STSdp), and peri-sylvian language (PSL). Data at https://balsa.wustl.edu/1B9VG and https://balsa.wustl.edu/5Xr71

The diffusion MRI is corrected for distortion and motion due to gradient nonlinearity, eddy current, motion, and B0 static field inhomogeneity using EDDY (Andersson and Sotiropoulos, 2016). The signal dropouts, susceptibility artefact, and their interaction with motion were also corrected (Andersson et al., 2018; Andersson et al., 2017). The resulting diffusion volumes are merged into a single volume and resampled in the subject’s real physical space aligned according to the ACPC convention. Diffusion modeling is performed using nerite orientation density imaging (NODDI) (Fukutomi et al., 2018; Zhang et al., 2012), and a Bayesian estimation of crossing fibers (Behrens et al., 2003; Sotiropoulos et al., 2016). Diffusion probabilistic tractography (Behrens et al., 2003) is also performed in a surface-based analysis (Donahue et al., 2016).

#### 2.5.3 Preliminary travelling subject data

Here, we show the preliminary results obtained from the initial TS data (as detailed in section 2.4). In the initial TS study (N=30), four healthy subjects participated and travelled across five sites and received MRI scanning with HARP in different scanners (4TS⨯5S), and twenty six subjects completed test-retest scans in any of 5 scanners (26TS⨯2/5S). Datasets were analyzed with the current version of preprocessing (see section 2.5.2) and each of the cortical thickness, myelin (non BC), and functional connectivity was parcellated using HCP MMP v1.0 (Glasser et al., 2016a) as described above (a part of the parcellated data in an exemplar subject [ID = 9503] was already shown in Figure 3 and 6). To investigate similarity of the data, each of the parcellated metrics was fed into an analysis of Spearman’s rank correlation across subjects and sites/scanners. Figure 7 shows the resultant similarity matrices which demonstrate higher correlation coefficients of within-subjects & between scanners than those of cross-subjects & between scanners in all the metrics of cortical thickness, myelin, and functional connectivity.

**Figure 7.**
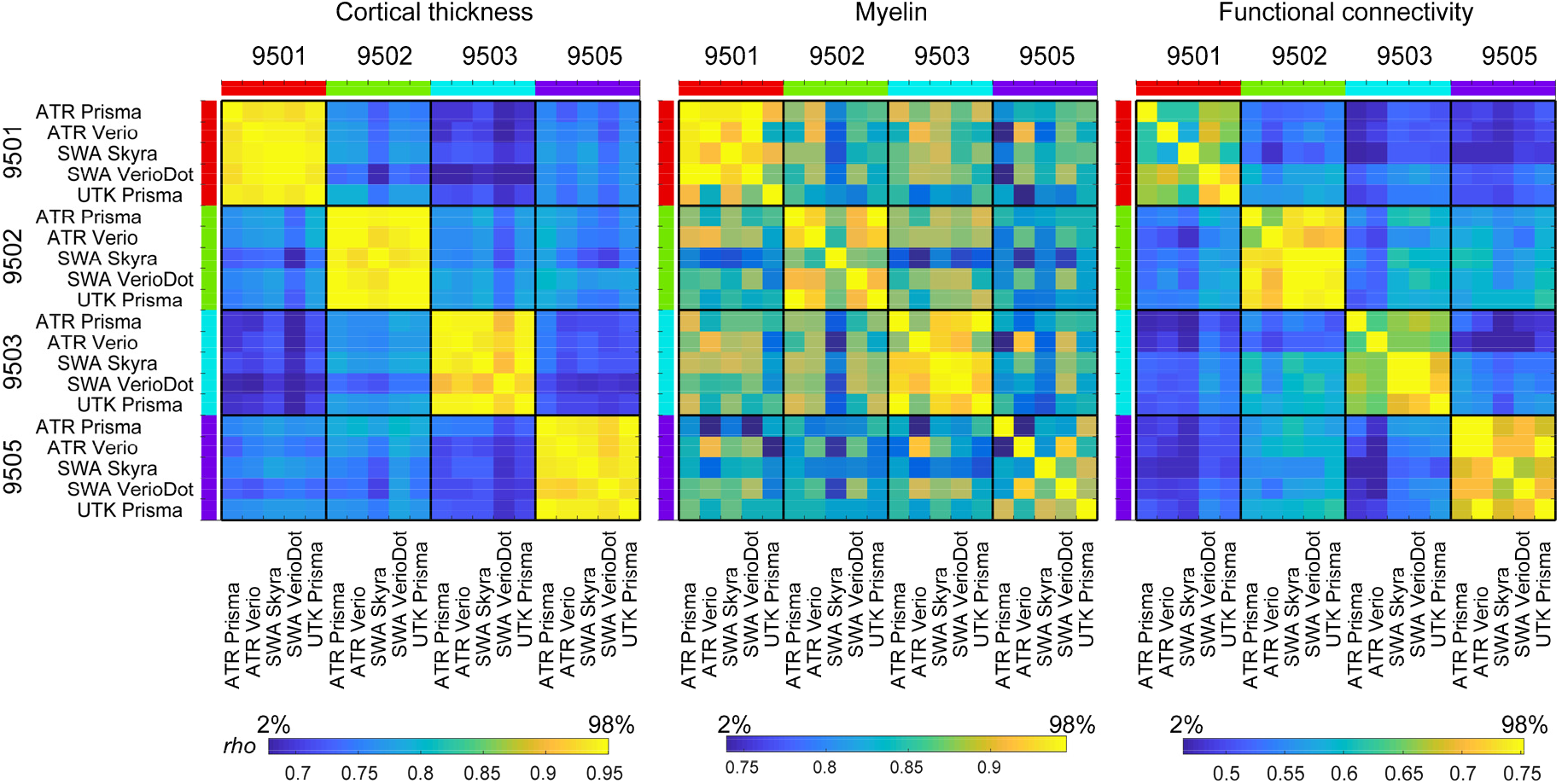
Similarity of the cortical metrics across subjects and sites/scanners in preliminary travelling subject study. From left to right show the correlation matrices of the parcellated cortical thickness, myelin (non BC) and functional connectivity in four travelling subjects (TS). Number of parcellated metrics used for analysis were 360 for thickness and myelin and 129,240 for functional connectivity, which cover all the cerebral cortex in both hemispheres. The correlation coefficient of Spearman’s rho is presented by a color bar placed at the bottom.

We also analyzed a different set of TS (N=26), who received test-retest scanning with the HARP protocol in the same MRI sites/scanners. The results (Fig. 8) showed greater similarity of cortical thickness, myelin map, and functional connectivity between test-retest data within subjects as compared with those with different subjects and/or scanners. The correlation coefficients of within-subject & within-scanner were again moderately high and comparable with those of within-subject & between-scanners in Fig 7.

**Figure 8.**
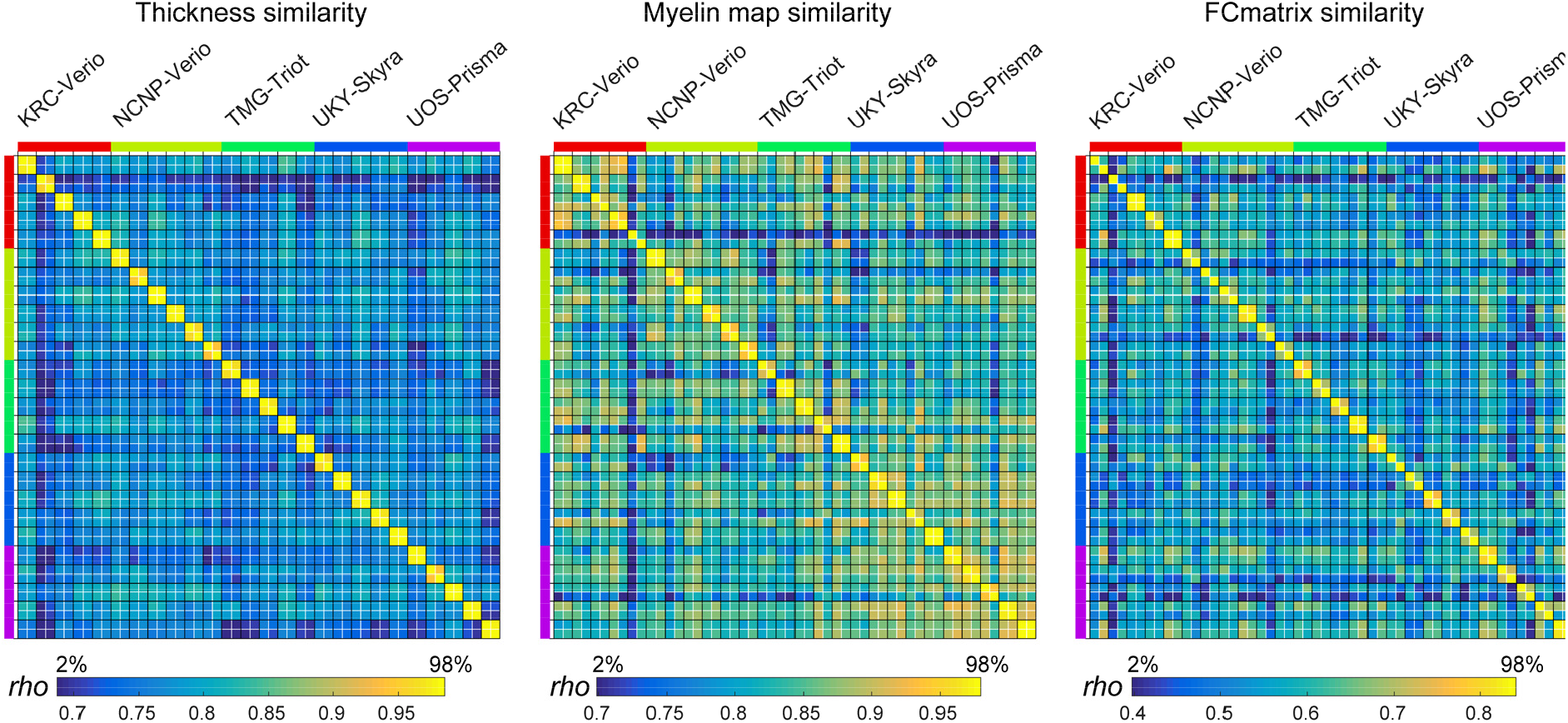
Test-retest results of cortical thickness, myelin (non BC) and resting-state functional connectivity (FC). Note that correlation adjacent to the diagonal and within each black box indicates a single subject’s test-retest correlation and is excellent in structure (thickness and myelin) and fairly good in FC. The different sites are colored along the left and top edges.

Table 4 summarizes the similarity values of all the TS30 data in Fig 7 and 8, classified into four types: within-subject & within-scanner, within-subject & between-scanner, between-subject & within-scanner, and between-subject & between-scanner. It is notable that the within-subject similarities are apparently higher than those of between-subject, indicating high sensitivity and reproducibility of subject-wise connectome. The between-subject similarities are smaller than within-subject and almost same across scanners, suggesting minimal bias between scanners and protocols. The within-subject & between-scanner similarity of the myelin map (0.89±0.05) was slightly degraded as compared with within-subject & within-scanner (0.95±0.03), suggesting the residual bias from transmit field across scans, for which we need to develop the correction method in future. That said, these preliminary datasets indicate that the HARP protocols and cortical parcellated analysis provide highly reproducible and specific pattern of subject-wise connectome, which may effectively enhance statistical harmonization (see Section 2.4) once the data was fully collected in this project.

**Table 4.** Summary of similarity of cortical measures in TS30.

| Type of correlation | Thickness | Myelin Map<br>(non BC) | FC matrix |
| --- | --- | --- | --- |
| Within-subject & within-scanner correlation (Subject N=26, combination N=26) | 0.97 (0.01) | 0.95 (0.03) | 0.72 (0.08) |
| Within-subject & between-scanner correlation (Subject N=4, combination N=40) | 0.94 (0.01) | 0.89 (0.05) | 0.69 (0.04) |
| Between-subject & within-scanner correlation (Subject N=30, combination N=250) | 0.76 (0.03) | 0.85 (0.05) | 0.55 (0.07) |
| Between-subject & between-scanner correlation (Subject N = 30, combination N=1200) | 0.76 (0.03) | 0.83 (0.05) | 0.55 (0.07) |

#### 2.5.4. Quality control

QC is implemented in several stages: 1) a brief image check during each scan; 2) an anomaly and abnormality inspection by the radiologists; 3) an assessment of raw data image quality when uploading data to the XNAT server; and 4) preprocessed image quality checks. QC 1 is conducted by site personnel and the participants are rescanned within the same session if scan time remains, if the images have major artifacts, such as those due to head movement. QC 2 is conducted by radiologists at the measurement site or other sites if any radiologist at the site is unable to check the images. QC 3 is manually conducted by researchers at the measurement sites before uploading the data to a server for all images in reference to the HCP QC manual (Marcus et al., 2013). After uploading the images to the XNAT servers, all images are first checked according to the DICOM file information as to whether the images are correctly updated. The researchers at each site are informed of missing DICOM files and any irregular parameters detected in the DICOM files. In QC 3, the T1w and T2w images are manually checked as to whether the face images are completely removed. Then, signal distributions of the myelin map are checked for outliers because of its sensitivity to several artifacts and errors such as motion, reconstruction of the images, and cortical surface reconstruction. Functional and diffusion images are automatically checked for outliers, and the images and data will be checked by visual inspection. In the QC 3 process, a QC pipeline will be implemented for checking the images (Marcus et al., 2013). QC 4 uses preprocessed CIFTI images that will be checked in several preprocessing steps. Any irregular scans and remarks are recorded in the clinical data servers and the information will be used when determining the eligibility criteria for each study.

### 2.6. Ethical regulation

Sharing neuropsychiatric patient data, which may contain information linked to subjects’ privacy, requires special attention (Sadato et al., 2019). Therefore, the Brain/MINDS Beyond project put NCNP as the core site for supporting ethical considerations. Before participating in the project, all institutions are required to receive approval from their ethical review board regarding their research plans. This includes the following points and ethical documentation: 1) MR images and clinical data of the participants may be shared within the Brain/MINDS Beyond project or Japanese/International scientific institutions for collaboration. De-identified MR images with limited clinical data (see below) may become publicly accessible on an open database for research purposes. 2) MR images of the participants may be compared with non-human primate MRI data. 3) Intellectual property rights originating from the research of the Brain/MINDS Beyond project shall be attributed to the institutes of the researchers and not the participants. All participants must provide written informed consent to participate in this project after receiving a complete explanation of the experiment.

The Japanese regulations for the sharing of personal information used for research purposes requires attention in dealing with two types of data: “individual identification codes” and “special care-required personal information” (http://www.japaneselawtranslation.go.jp/law/detail/?id=2781&vm=04&re=01). Individual identification codes are direct identifiers—information sufficient to identify a specific individual. Special care-required personal information represents indirect identifiers needing special care in handling so as not to cause potential disadvantages to participants. In consideration of these regulations, data accompanied with the MR images are limited in the publicly accessible open database, and only include 5-year age bins, sex, diagnostic information, handedness, simple socioeconomic status, clinical scale scores, and sleepiness scale scores. In the Brain/MINDS Beyond project, we exclude the datasets of MR images containing facial information from the data in the publicly accessible open database.

### 2.7. Data sharing

In the current provisional plan of sharing the collected data, we have designated three types of data sharing:

1. Access via an open database: de-identified MR images and limited clinical data are to become publicly accessible for research purposes after the research period ends. The initial release will be scheduled in 2024. Basic demographic and clinical characteristics such as 5-year age bin, sex, socioeconomic status, (premorbid) estimated intellectual quotient, main diagnosis, representative scale scores for each disease and sleepiness during rsfMRI scan will be shared.
2. Application-based sharing: MR images and the clinical datasets are shared after receiving application approval for data usage by the Brain/MINDS Beyond human brain MRI study working group. Applicants are required to obtain approval of their research plan from the ethical review board of their institution and request the dataset type in the application form. The working group discusses the eligibility of the applicants, as well as the availability of the requested dataset, the ethical consideration in the Brain/MINDS Beyond site(s), and any conflict from other applications. Data is released from the distribution server of the Brain/MINDS Beyond project with limited access.
3. Collaboration-based sharing: This form of sharing is used for individual collaborative studies. A research proposal collaborating with the institute(s) in the Brain/MINDS Beyond project is approved by the ethical review board of the institute(s). Data is shared from the relevant institute(s).

## 3. Discussion

The Brain/MINDS Beyond human brain MRI study expands upon research from previous multi-site neuroimaging studies in Japan and provides high quality brain images by standardizing multiple MRI scanners and protocols. An unbiased and quantitative assessment of cortical structure and function may be needed for sensitive and specific predictions of any dynamics, perturbations, or disorders of the brain system. Multi-modal cross-disease image datasets are systematically and properly acquired, analyzed, and shared to enable investigation of common and disease-specific features for psychiatric and neurological disorders with a high sensitivity and specificity. A distinct feature of this project is to include a study design with the TS project, which enables harmonizing the multi-site data from lower (i.e. preprocessing) to higher levels (i.e. statistics). The harmonization protocols are available at http://mriportal.umin.jp. The Brain/MINDS Beyond human brain MRI project can provide brain imaging biomarkers that are applicable to therapeutic targets and diagnostic supports.

To date, several national projects have applied high-quality multimodal MRI protocols, in addition to a preprocessing pipeline, to a large cohort (e.g., HCP, UK biobank, and ABCD). Unlike these multi-site projects, we plan to investigate brain organization associated with brain disorders that occur throughout the lifespan and to develop imaging biomarkers that can be implemented in clinical trials. To facilitate the collection of a larger number of patients with different brain disorders, multiple clinical research sites are participating in this project and cooperating for standardized data acquisitions. The core of the project began from establishing a standardized protocol (i.e. HARP) based on five 3T MRI scanners, but it will continue to develop a comparable protocol for other types of scanners/vendors. The protocol is designed not only for high-resolution structural MRI and high-quality resting-state fMRI, but also for diffusion MRI and other imaging—including scans for correcting distortions. The preprocessing is performed with a surface-based multi-modal analysis to minimize bias largely generated from the variability in cortical folding across subjects (Coalson et al., 2018; Glasser et al., 2016b). The preliminary data demonstrated high quality MRI images and the fidelity of structural and functional brain organizations across scanners/sites. The signal-to-noise ratio of MRI images was very high across scanners/sites (Figure 3A). The cortical metrics of structure (myelin map, thickness) (Figure 3B-C) were comparable to those previously reported in the literature (Fischl and Dale, 2000; Glasser and Van Essen, 2011), as well as the functional connectivity related to eye movements involving FEF and PEF (Figure 6A) (Amiez and Petrides, 2009) and a language network involving left 55b, 44/45, STS, and PSL (Figure 6B) (Glasser et al., 2016a). These findings suggest that a surface-based parcellated analysis may provide useful and reliable metrics concerning cortical structure, function, and connectivity, and may potentially contribute to the establishment of multi-modal imaging biomarkers of brain disorders. The initial trial with 30 TS also demonstrated the highly reproducible and specific pattern of subject-wise connectome across five scanners, suggesting the reliability of our prospective harmonization (e.g. protocols and preprocessing) and promising future retrospective (i.e. statistical) harmonization. The residual bias of myelin map is presumably due to differences of transmit field across scans and needs to be corrected in future preprocessing pipeline.

The TS approach is a novel harmonization method for multi-site brain image data (Yamashita et al., 2019), which has proven that measurement bias from MRI equipment and protocols can be differentiated from sampling bias between sites. Instead of using a previously applied GLM, we plan to expand the statistical approach to a GLMM in this project. One of the obstacles of the GLMM approach is that it requires a larger number of total scans compared to those in a GLM approach; overlapping scans at hub sites are required for all TS participants to ensure the data connectivity; additionally, a larger number of TS participants is required in the TS project because the degree of freedom can be reduced in the GLMM. However, one of the benefits of the GLMM approach includes that it is flexible with the variability in data acquisition—such as the number of scans per participant and length of scan time per protocol; thus, is suitable for a big project. Furthermore, this approach allows the addition of another site, scanner, and protocol to an existing TS network, which can deal with the future upgrades of scanners and protocols. In fact, the scanners at two sites (UHI and SWA) were upgraded to a MAGNETOM Skyra fit (Siemens Healthcare GmbH, Erlangen, Germany) for institutional reasons after the Brain/MINDS Beyond project had started. Therefore, we customized the TS for two sites to ensure that the data are properly connected before and after the upgrades. Also, the project welcomes other sites to participate in the TS network.

Because this project focuses on various brain disorders across the lifespan, we aim to identify common and disease-specific features of psychiatric and neurological disorders. While some case-control studies suggest possible neural mechanisms in a psychiatric disease, other studies suggest that the effects may not be specific to a single entity but instead may be shared across multiple neuropsychiatric disorders (Hibar et al., 2018; Schmaal et al., 2017; Schmaal et al., 2016; van Erp et al., 2016). Such non-specificity may be at least partly addressed by investigating diseases across the lifespan, since some of brain changes reported in psychiatric disorders also occur in aging or development in healthy subjects, e.g. volumetric changes in subcortical structures in schizophrenia (Okada et al., 2016; van Erp et al., 2016) and in healthy aging (O’Shea et al., 2016; Wang et al., 2019). We initially coordinated with 13 sites to explore various psychiatric and neurological disorders throughout the lifespan and to make use of a powerful harmonization method. Therefore, this project aims to identify both the common and disease-specific pathophysiology features of psychiatric and neurological disorders, which will hopefully lead to imaging biomarkers for general clinical practice and the development of candidate therapeutic targets for future clinical trials.

We established task fMRI scans using EMOTION and CARIT in HARP to evaluate the validity, reliability and applicability of harmonization. The result of task fMRI may be used for validation of the resting-state fMRI based cortical surface registration, and parcellation. The TS project performing task fMRI is now under planning and hopefully will be completed in coming years.

In conclusion, the Brain/MINDS Beyond human brain MRI project began with the participation of 13 clinical research sites—all of which have setup brain image scans using the standard MRI scanners and protocols, conducted TS scans, and will share acquired data with the project and the public in the future, and commit to the analysis and publication of the data. To the best of our knowledge, this is the first human brain MRI project to explore psychiatric and neurological disorders across the lifespan. The project aims to discover robust findings which may be directly related to the common or disease-specific pathophysiology features of such diseases and facilitate the development of candidate biomarkers for clinical application and drug discovery.

## Abbreviations

DALYs: disability-adjusted life years
MRI: magnetic resonance imaging
HCP: Human Connectome Project
ABCD: Adolescent Brain Cognitive Development
BPD: bipolar disorder
MDD: major depressive disorder
DecNef: Decoded Neurofeedback
ASD: autism spectrum disorder
ADNI: Alzheimer’s Disease Neuroimaging Initiative
AD: Alzheimer’s disease
MCI: mild cognitive impairment
PPMI: Parkinson’s Progression Markers Initiative
PD: Parkinson’s disease
T1w: T1-weighted
T2w: T2-weighted
rsfMRI: resting state functional MRI
CRHD: Connectome Related to Human Disease
GLM: general linear model
TS: traveling subject
AMED: Japan Agency for Medical Research and Development
Brain/MINDS Beyond: Strategic International Brain Science Research Promotion Program
HARP: Harmonization protocol
DWI: diffusion-weighted imaging
QC: quality control
MNI: Montreal Neurological Institute
MSM: multi-modal surface matching
GLMM: general linear mixed model
CIFTI: Connectivity Informatics Technology Initiative
FEF: frontal eye field
PEF: premotor eye field
PSL: peri-sylvian language
STS: superior temporal sulcus
NODDI: nerite orientation and density imaging

## Acknowledgment

This research was supported by the Agency for Medical Research and Development (AMED) under Grant Numbers JP20dm0307001 (K.K.), JP20dm0307002 (Y.O.), JP20dm0307003 (T.Han.), JP20dm0307004 (K.K.), JP20dm0307008 (M.K.), JP20dm’0307020 (N.S.), JP20dm0207069 (S.K.) and JP20dm0307006 (T.Hay.). This study was also supported by the University of Tokyo Center for Integrative Science of Human Behaviour (CiSHuB) and the International Research Center for Neurointelligence (WPI-IRCN) at the University of Tokyo Institutes for Advanced Study (UTIAS).

## Author Contributions

Shinsuke Koike, Conceptualization, Project administration, Funding acquisition, Writing – original draft, Software, Resources.

Saori C Tanaka, Conceptualization, Project administration, Data Curation, Software, Resources, Writing – original draft.

Tomohisa Okada, Methodology, Investigation.

Toshihiko Aso, Investigation, Formal Analysis, Validation.

Michiko Asano, Investigation, Data curation.

Norihide Maikusa, Methodology, Resources.

Kentaro Morita, Writing – original draft.

Naohiro Okada, Investigation, Writing – original draft.

Masaki Fukunaga, Methodology.

Akiko Uematsu, Methodology, Investigation.

Hiroki Togo, Methodology, Investigation.

Atsushi Miyazaki, Methodology, Investigation.

Katsutoshi Murata, Methodology.

Yuta Urushibata, Methodology.

Joonas Autio, Methodology.

Takayuki Ose, Methodology.

Junichiro Yoshimoto, Methodology.

Toshiyuki Araki, Writing – original draft.

Matthew F Glasser, Software, Writing - reviewing and editing.

David C Van Essen, Writing – reviewing and editing.

Megumi Maruyma, Project administration.

Norihiro Sadato, Investigation, Funding acquisition, Project administration.

Mitsuo Kawato, Conceptualization, Funding acquisition, Project administration.

Kiyoto Kasai, Investigation, Funding acquisition, Project administration, Supervision.

Yasumasa Okamoto, Investigation, Funding acquisition.

Takashi Hanakawa, Project administration, Funding Acquisition, Methodology, Investigation, Resources, Writing – original draft.

Takuya Hayashi, Conceptualization, Project administration, Funding Acquisition, Software, Resources, Formal Analysis, Writing - original draft, reviewing and editing.

## Conflict of interest

Katsutoshi Murata and Yuta Urushibara are employed by Siemens Healthcare K.K., Tokyo, Japan. The other authors report no financial relationships with commercial interests.

## Data availability

The data presented in Figure 3 and 6 are available at BALSA (https://balsa.wustl.edu/study/show/npD26). Harmonization protocols and other information of the project are available at the BrainMINDS beyond MRI portal site (http://mriportal.umin.jp/?lang=en). The tool for DICOM to NIFTI conversion, folder structure, derivation of imaging parameters is available at https://github.com/RIKEN-BCIL/BCILDCMCONVERT, See also Data sharing section in details of data obtained in future in this project. For proposal and requests for the data usage, please contact to Saori Tanaka.

## Notes

### Summary of Updates

In this revision, we carried out additional analyses of test-retest data (N=26), as shown in Fig 8. We also replaced myelin maps used in Fig 3, 7 and 8 to those of non biasfield corrected to clarify the need for future elaboration of preprocessing. Overall, the preliminary test-retest in travelling subjects (N=30) not only demonstrated moderately high reliability and specificity to subject-wise connectome but also an issue to be solved as for correction of transmit field biasfield in future.

