## Supplemental Table 1 for "Brain/MINDS Beyond Human Brain MRI Project: A Protocol for Multi-Site Harmonization across Brain Disorders Throughout the Lifespan"

Running Head: Brain/MINDS Beyond MRI study

Shinsuke Koike, M.D., Ph.D.[1,2,3,4]; Saori C Tanaka, Ph.D.[5]; Tomohisa Okada, M.D., Ph.D.[6]; Toshihiko Aso, M.D., Ph.D.[7]; Michiko Asano, Ph.D.[1]; Norihide Maikusa, Ph.D.[1,8]; Kentaro Morita, M.D., Ph.D.[9]; Naohiro Okada, M.D., Ph.D.[2,4,10]; Masaki Fukunaga, Ph.D.[11]; Akiko Uematsu, Ph.D.[1]; Hiroki Togo, MSc.[8]; Atsushi Miyazaki, Ph.D.[12]; Katsutoshi Murata, MSc.[13]; Yuta Urushibata, MSc.[13]; Joonas Autio, Ph.D.[7]; Takayuki Ose, MSc[7]; Junichiro Yoshimoto, Ph.D.[5]; Toshiyuki Araki, M.D., Ph.D.[14]; Matthew F Glasser, M.D., Ph.D.[15,16,17]; David C Van Essen, Ph.D.[15]; Megumi Maruyama, Ph.D.[18]; Norihiro Sadato, M.D., Ph.D.[11]; Mitsuo Kawato, Ph.D.[5,19]; Kiyoto Kasai, M.D., Ph.D.[2,3,4,10]; Yasumasa Okamoto, M.D., Ph.D.[20]; Takashi Hanakawa, M.D., Ph.D.[8,21]; Takuya Hayashi, M.D., Ph.D.[7]; Brain/MINDS Beyond Human Brain MRI Group

#### Supplementary Materials

### Table of Contents

**Supplementary Table S1. Main imaging parameters of structural (T1_MPR and T2_SPC), functional MRI (fMRI) and diffusion weighted MRI (DWI) scans in HARP protocol.** ........................................ 2

**Supplementary Table S2. Traveling subject project protocol.** .................................................. 4

**Supplementary Table S1. Main imaging parameters of structural (T1_MPR and T2_SPC), functional MRI (fMRI) and diffusion weighted MRI (DWI) scans in HARP protocol.**

| Scanner (System version) | | Prisma (VE11C) | Skyra (VE11C) | Verio (VD13A) | Verio (VB17A) | Trio (VB19A) |
|---|---|---|---|---|---|---|
| **T1_MPR** | | | | | | |
| | FOV(SI × AP × RL) [mm] | 256 × 240 × 179.2 | | | | |
| | Orientation | sagittal (PE dir. : A >> P) | | | | |
| | Matrix size | 320 × 300 × 224 | | | | |
| | Slice oversampling [%] | 7.1 | | | | |
| | Resolution [mm] | 0.8 × 0.8 × 0.8 | | | | |
| | TR/TE/TI [msec] | 2500/2.18/1000 | | | | |
| | Scan time [min:sec] | 5:22 | | | | |
| | Flip angle [deg] | 8 | | | | |
| | Acceleration | Phase partial Fourier: 6/8, GRAPPA (factor:2, ref. line:32) | | | | |
| | Bandwidth [Hz/Px] | 220 | | | | 210 |
| | Echo spacing [msec] | 7.9 | | | | 7.8 |
| | Sampling Spacing [μsec] | 7.1 | | | | |
| | Fat Sat | Water excit. fast | | | | |
| | Filter | Distortion Corr OFF, Prescan Normalize ON | | | | |
| **T2_SPC** | | | | | | |
| | FOV(SI × AP × RL) [mm] | 256 × 240 × 179.2 | | | | |
| | Orientation | sagittal (PE dir. : A >> P) | | | | |
| | Matrix size | 320 × 300 × 224 | | | | |
| | Resolution [mm] | 0.8 × 0.8 × 0.8 | | | | |
| | Slice oversampling [%] | 7.1 | | | | |
| | TR/TE [msec] | 3200/564 | | 3200/565 | 3200/564 | 3200/562 |
| | Scan time [min:sec] | 5:31 | | 5:22 | 6:26 | |
| | Excitation | variable flip angles (Flip angle mode: T2 var) | | | | |
| | Turbo factor | 314 | | 326 | 167 (Slice turbo factor 2) | |
| | Acceleration | GRAPPA (factor:2, ref. line:32) | | | | |
| | Bandwidth [Hz/Px] | 744 | 679 | | | 781 |
| | Echo spacing [msec] | 3.86 | | 3.72 | | 3.48 |
| | Sampling Spacing [μsec] | 2.1 | 2.3 | | | 2.1 |
| | Filter | Distortion Corr OFF, Prescan Normalize ON, Image Filter (Sharp: Edge Enhanvement 3 & Smoothing 3) | | | | |
| **fMRI BOLD** | | | | | | |
| | FOV(RL × AP × SI) [mm] | 206 × 206 × 144 | | | | |
| | Orientation | transverse (PE dir. : swapped alternatively btw A >> P & P >> A) | | | | |
| | Matrix size / Slices | 86 × 86 / 60 | | | | |
| | Resolution [mm] | 2.4 × 2.4 × 2.4 | | | | |
| | TR/TE [msec] | 800/34.4 | | | | |
| | #measurements | 375 | | | | |
| | Scan time [min:sec] | 5:08 | | | | |
| | Flip angle [deg] | 52 | | | | |
| | Acceleration | Partial Fourier: OFF, Multi-band (factor:6) | | | | |
| | Bandwidth [Hz/P × ] | 2076 | | | | 2326 |
| | Echo spacing [msec] | 0.63 | | | | 0.57 |
| | Filter | Prescan Normalize ON | | | | |
| **fMRI Spin-echo fieldmap** | | | | | | |
| | FOV(RL × AP × SI) [mm] | 206 × 206 × 144 | | | | |
| | Orientation | transverse (PE dir. : swapped alternatively btw A >> P & P >> A) | | | | |
| | Matrix size | 86 × 86 × 60 | | | | |
| | Resolution [mm] | 2.4 × 2.4 × 2.4 | | | | |

| | | | | | |
|---|---|---|---|---|---|
| | TR/TE [msec] | 6100/60 | | | |
| | Scan time [min:sec] | 0:06 | | | |
| | Flip angle [deg] | 90/180 | | | |
| | Acceleration | Phase Partial Fourier OFF | | | |
| | Bandwidth [Hz/P × ] | 2076 | | 2154 | 2326 |
| | Echo spacing [msec] | 0.63 | | | 0.57 |
| | Filter | Prescan Normalize ON | | | |
| **DWI** | | | | | |
| | FOV(RL × AP × SI) [mm] | 204 × 204 × 144 | | | |
| | Orientation | transverse (PE dir. : swapped alternatively btw A >> P & P >> A) | | | |
| | Matrix size | 120 × 120 × 84 | | | |
| | Resolution [mm] | 1.7 × 1.7 × 1.7 | | | |
| | TR/TE [msec] | 3600/79.0 | 3600/89.0 | | 3600/94.0 |
| | Scan time [min:sec] | 3:29 (AP)<br>3:32 (PA) | 4:50 (AP)<br>4:54 (PA) | | |
| | Flip angle [deg] | 90/180 | | | |
| | Acceleration | Partial Fourier: 6/8, Multi-band [factor: 3] | Partial Fourier: 6/8, GRAPPA (ref line 36) Multi-band [factor: 3] | | |
| | Bandwidth [Hz/Px] | 1984 | 1544 | 1436 | 1736 |
| | Echo spacing [msec] | 0.62 | 0.74 | 0.78 | 0.70 |
| | Filter | Prescan Normalize ON | | | |
| | b-values [s/mm$^2$] | 0/700/2000 | | | |
| | Diffusion direction | 5/16/32 (AP)<br>6/16/32(PA) | 7/20/40(AP)<br>8/20/40(PA) | | |
| | Diffusion scheme | Monopolar | | | |

**Supplementary Table S2. Traveling subject project protocol.**

| Site | Protocol for target population | Additional protocol for TS | HARP additional measurement for TS | | | Other TS measurement sites[a] | | Total number of CRHD and HARP measurements |
|---|---|---|---|---|---|---|---|---|
| | | | 10-min additional rsfMRI | QSM | ASL | Hub site | | |
| **UTK** | CRHD | HARP, SRPB (Prisma 64-ch head coil) | ✓ | ✓ | ✓ | UTI | UHI | 7 |
| **UTI** | CRHD | HARP, SRPB (GE MR750W) | ✓ | ✓ | ✓ | UTK, ATR | | 8 |
| **ATR** | CRHD | HARP (Prisma, Verio), SRPB (Prisma, Verio) | ✓ | ✓ | ✓ | UTK | SWA[2] | 8 |
| **FUM** | HARP | NA | ✓ | ✓ | ✓ | UTK | SWA, NCNP | 6 |
| **TMG** | HARP | SRPB | ✓ | | | UTK | SWA, UHI | 6 |
| **SWA** | HARP | NA | ✓ | ✓ | ✓ | UTK, UTI | | 6 |
| | HARP | SRPB (Verio)[b] | ✓ | ✓ | ✓ | UTI | | 5 |
| **NCNP** | HARP | SRPB | ✓ | | ✓ | UTK | JTD, FUM | 6 |
| **JTD** | HARP | CRHD | ✓ | | | UTK | NCNP, UKY | 6 |
| **UOS** | HARP | CRHD, SRPB (Trio Tim) | ✓ | ✓ | ✓ | ATR | UKY, TMG | 6 |
| **UHI** | HARP | SRPB (Verio)[c] | ✓ | ✓ | ✓ | UTK | NCNP, SWA | 6 |
| **UNG** | HARP | NA | ✓ | | | UTK | TMG, NCNP | 6 |
| **UKY** | HARP | NA | ✓ | ✓ | ✓ | ATR | UOS, NCNP, JTD | 6 |
| **KRC** | HARP | NA | ✓ | | | ATR | UKY, UNG | 6 |

Abbreviations: TS, traveling subject; rsfMRI, resting state functional MRI; QSM, quantitative susceptibility mapping; ASL, arterial spin labeling; UTK, The University of Tokyo ECS (Komaba Campus); UTI, The University of Tokyo IRCN; FUM, Fukushima Medical University; TMG, Tamagawa Academy & University; SWA, Showa University; NCNP, National Center of Neurology and Psychiatry; JTD, Juntendo Hospital; ATR, Advanced Telecommunications Research Institute International; UOS, Osaka University; UHI, Hiroshima University; UNG, Nagoya University; UKY, Kyoto University; KRC, Kyoto University Kokoro Reserch Center; IR, Innovative Research Group in Brain/MINDS Beyond; BM, Brain/MINDS project; CRHD, Human Connectome Studies Related To Human Disease protocol; HARP, HARmonized protocol; SRPB, Strategic Research Program for Brain protocol (for previous multicenter study projects).

a In the hub site(s), participants were measured using Prisma CRHD and HARP protocols. We planned the measurements in other sites per site, considering the feasibility of measurement (project, machine, location, availability, etc.) and the mixture of the pairs between measurements.
b The participants who were measured using Verio SRPB in Showa University before a machine upgrade participated in the measurements.
c Some of the participants were measured using Verio SRPB in Hiroshima University, and Prisma CRHD and HARP in Advanced Telecommunications Research Institute International since the machine in Hiroshima University had been upgraded before the project started.
